# Function and evolution of B-Raf loop dynamics relevant to cancer recurrence under drug inhibition

**DOI:** 10.1101/2020.01.13.904052

**Authors:** Gregory A. Babbitt, Miranda L. Lynch, Matthew McCoy, Ernest P. Fokoue, André O. Hudson

## Abstract

Oncogenic mutations in the kinase domain of the B-Raf protein have long been associated with cancers involving the MAPK pathway. One constitutive MAPK activating mutation in B-Raf, the V600E (valine to glutamate) replacement occurring adjacent to a site of threonine phosphorylation (T599) occurs in many types of cancer, and in a large percentage of certain cancers, such as melanoma. Because ATP binding activity and the V600E mutation are both known to alter the physical behavior of the activation loop in the B-Raf ATP binding domain, this system is especially amenable to comparative analyses of molecular dynamics simulations modeling various genetic and drug class variants. Here, we employ machine learning enabled identification of functionally conserved protein dynamics to compare how the binding interactions of four B-Raf inhibitors impact the functional loop dynamics controlling ATP activation. We demonstrate that drug development targeting B-Raf has progressively moved towards ATP competitive inhibitors that demonstrate less tendency to mimic the functionally conserved dynamic changes associated with ATP activation and leading to the side effect of hyperactivation (i.e. inducing MAPK activation in non-tumorous cells in the absence of secondary mutation). We compare the functional dynamic impacts of V600E and other sensitizing and drug resistance causing mutations in the regulatory loops of B-Raf, confirming sites of low mutational tolerance in these regions. Lastly, we investigate V600E sensitivity of B-Raf loop dynamics in an evolutionary context, demonstrating that while sensitivity has an ancient origin with primitive eukaryotes, it was also secondarily increased during early jawed vertebrate evolution.

## Introduction

From both a patient’s and doctor’s perspective, perhaps the most dreaded aspect of many cancers are their ability to undergo recurrence following therapeutic intervention. The recurrence of cancer can have obvious negative impact on patient psychology (Andersen et al., 2005; Yang et al., 2008), doctor-patient relations (Janz et al., 2017) and questions the very concept of a ‘cure’ for cancer. The molecular mechanisms and somatic evolutionary processes that underlie the phenomena of recurrence are complex and do not act singularly within any given tumor, thereby greatly exacerbating the clinical difficulty of treatment and post-treatment monitoring. There are currently three molecular mechanistic frameworks for potentially explaining cancer recurrence. The traditional view of ‘mutational deregulation’ invokes a progressive molecular evolutionary Fisherian or ‘runaway’ process whereby tumor progression begins with one or several ‘driver mutations’ that start a cascade of secondary mutation and subsequent deregulation of growth and proliferation followed by eventual chromosomal disruption through aneuploidy and chromothrypsis (Brown et al., 2019). Under this conceptual framework, cancer progression is largely due to a mutational process that progressively deregulates cell proliferation and frames cancer recurrence as a problem analogous to a ‘whack a mole’ game that gets more difficult as the game progresses. A second molecular evolutionary perspective that has gained recent popularity invokes Darwinian natural selection at the somatic level of the tumor microenvironment (Gatenby et al., 2019; Zhang et al., 2017a). Under this view, the cancer treatment regime, in creating a local tumor microenvironment that is selectively detrimental to the survival and proliferation of cancer cells, is thought to create a selective microenvironment that favors more robust and treatment-resistant cancers that can recur if therapy is not modulated accordingly. Cancer recurrence under this new paradigm is framed as a game of strategy like chess, where the competitive environment changes fluidly with the progression of the game. Both of these conceptual frameworks for our understanding of cancer recurrence rely upon disruption of mutation-selection balance at a tissue level (Bürger, 1989; Hartl, 1977) acting through the complementary mechanisms of somatic mutation and apoptosis (Nunney, 2003).

A third and less common cause of cancer recurrence, known as the ‘hyperactivation paradox’ does not rely upon disruption of selection-mutation balance and can be invoked to explain cancer recurrence in the absence of secondary mutations. Hyperactivation is observed when some cancer inhibitors have opposing effects in tumor and normal wildtype cells, repressing growth pathways in tumor tissues while paradoxically activating them in normal healthy tissue (Hatzivassiliou et al., 2010; Poulikakos et al., 2010; Villanueva et al., 2011). The mechanism of hyperactivation may be less of a paradox when one considers that small molecule inhibitors are often targeting the binding sites of natural agonists (i.e. signal pathway activators) in both tumor and wildtype cells. In growth activated tumor cells, these drugs will competitively bind the target protein, thus blocking the functional role of the natural agonist in the pathway. However, in growth deactivated cells, these drugs might mimic certain functional aspects of the agonist, allowing partial activation of growth pathways in the presence of the drugs. While not a ‘game’ that most would choose to play, this effect of functional mimicry in some drugs might be analogous to ‘poking a sleeping bear’, where cancer recurrence can be caused in the absence of secondary mutation by a drug’s unintended signal activation of the growth pathway in nearby normal cells.

Proteins involved in growth signaling pathways, such as mitogen-activated protein kinases (MAPK), are often very labile in structure, involving highly dynamic regions of intrinsic disorder (e.g. activation loops) that facilitate complex dynamic shifts between functional states that ultimately govern interactions with other proteins on the pathway (Montagut and Settleman, 2009; Wan et al., 2004). Mutations that potentially alter function in highly dynamic loop regions are often implicated in many cancers. A well-studied oncogenic system exists in the ATP binding domain of the B-Raf protein, a serine/threonine-protein kinase that activates the MAPK (i.e. RAF-MEK-ERK) growth pathway (Figure 1). B-Raf is a member of the Raf family of serine/threonine kinases, along with A-Raf and C-Raf, and is the member with the highest basal activity of the three (Roskoski, 2010). B-Raf is a 766 amino acid protein with gene on chromosome 7 (7q34). There are 3 primary conserved domains for B-Raf, CR1, 2, and 3. CR3 (residues 443-735) is the domain that contains the enzymatic ATP kinase domain (KD, residues 457-714). B-Raf regulation functions primarily by the changes in the position and dynamics in several loop regions upon interaction of ATP with the hydrophobic binding pocket formed by V471, A481, L514, T529, W531, and C532. Within the KD, the glycine-rich P-loop (ATP-phosphate stabilizing loop) spans residues 464-471 and functions to anchor ATP during enzymatic activity. The activation segment or activation loop (residues 594-623) within the KD is partially disordered and contains the serine and threonine phosphorylation sites (S602 and T599). Thus, the loop regions primarily affected by ATP binding interaction are the ATP phosphate stabilizing P-loop at residues 464-471, the catalytic loop at residues 574-581 that captures and transfers the ATP γ-phosphate, and the activation loop segment and DFG motif at residues 594-623 which breaks hydrophobic interactions with the P-loop (i.e. activating the KD) with negative charge during phosphorylation of T599. Activating mutations in the KD can often lead to disrupted conformation and constitutive ERK activation and many mutations in this region are associated with loss of regulatory function and/or abnormal dimerization (Durrant and Morrison, 2018). In particular, a V600E mutation in the activation loop causing the constitutive activation of the B-Raf signaling is one of the most common mutations in many type of cancer, as well as a high percentage of certain cancers like melanoma. B-Raf mutants appear in multiple cancers, notably in melanoma, and colorectal, but also less frequently in non-small cell lung cancer, breast, ovarian, and several others (Davies et al., 2002; Rezaei Adariani et al., 2018; Wan et al., 2004). Greater than 50% melanomas harbor B-Raf activating mutations, and of these 90% involve the three bases of codon 600, and of these 90% are the single nucleotide missense mutation at position 1799 resulting in V600E substitution (although V600 K, D, R have also been less commonly observed (Ascierto et al., 2012)).

**Figure 1.**
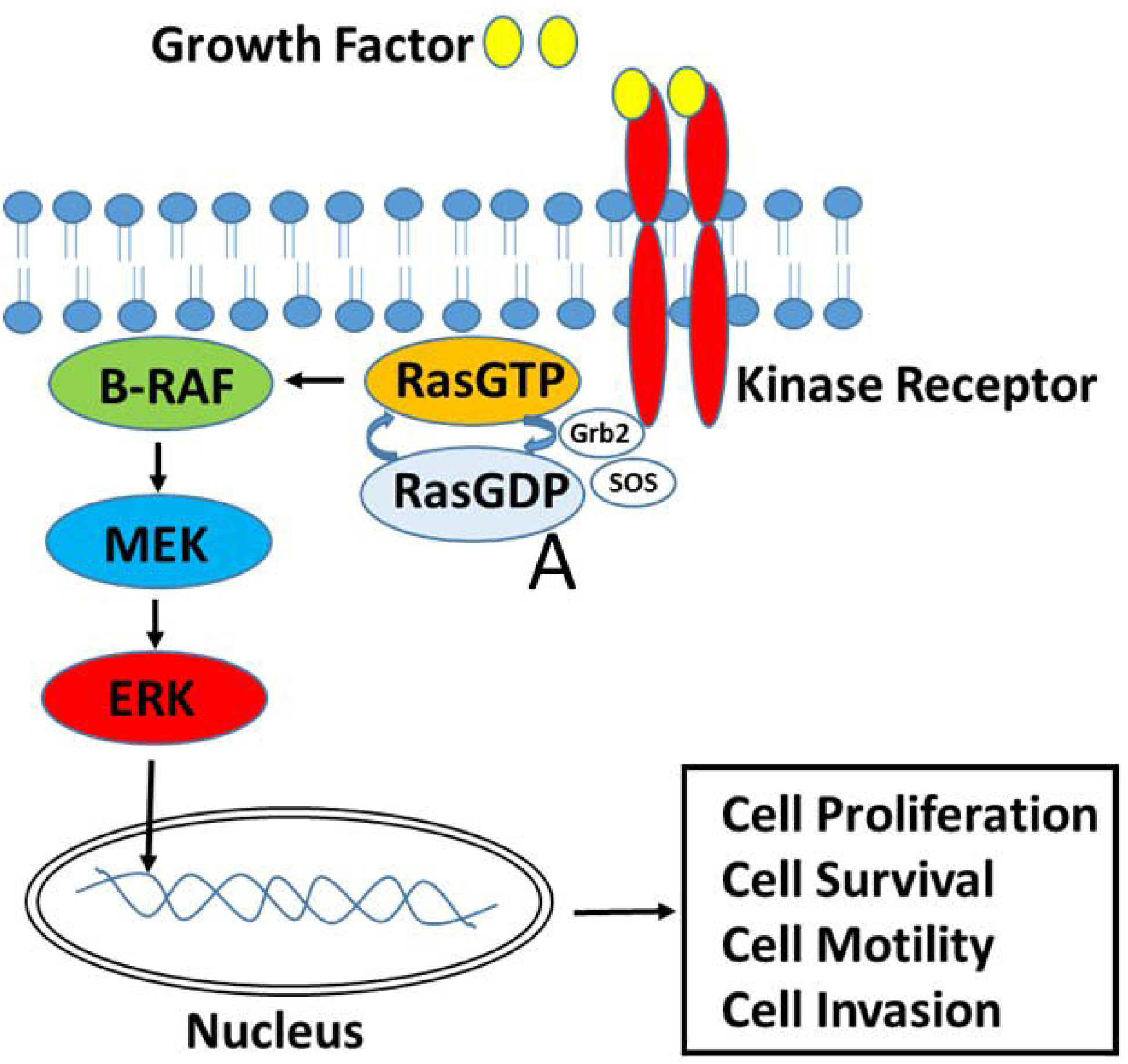
Overview of role of BRAF in mitogen-activated protein kinase pathway (MAPK) pathway. Growth factor/s binds to the receptor (EGFR) on the cell surface which leads to phosphorylation and subsequent activation. The scaffold proteins (GRB2 and SOS) facilitate the removal of GDP from RAS. RAS then binds GTP which leads to activation. Activated RAS undergoes a conformational change which leads to the phosphorylation and activation of BRAF, MEK and ERK. Activated ERK is shuttled to the nucleus where it is involved in recruiting transcription factors involved in the up regulation of genes involved in cell proliferation, cell survival, cell motility and cell invasion. The BRAF V600E mutation allows for constitutive activation of BRAF to facilitate downstream signaling through MEK and ERK circumventing upstream activation/regulation.

Several first generation B-Raf inhibitors designed to target V600E mutants cells have been recently discovered to induce hyperactivation in normal wildtype cells in the absence of V600E (Hatzivassiliou et al., 2010; Poulikakos et al., 2010; Villanueva et al., 2011). Now, a new generation of B-Raf inhibitors that break the hyperactivation paradox are currently under development and clinical trial (Zhang et al., 2015a). Raf inhibitors are generally designed to occupy the ATP binding pocket preventing phosphorylation. Why this region suffers from such low mutational tolerance, and why some Raf inhibitors prove vulnerable to hyperactivation while others do not, are currently open questions. A comparative analysis of the all-atom molecular dynamics (MD) of the B-Raf KD in both its wildtype and various mutant states during interaction with small molecule variants of this drug class could prove illuminating with regard to the problems of B-Raf hyperactivation and mutational tolerance more generally. Three B-Raf inhibitors are currently in use. These include sorafenib, a potent hyperactivation inducer in normal cells, (Zhang et al., 2015b, 2017b) as well as dabrafenib and vemurafenib, two more modern inhibitors that also show potential problems with hyperactivation (Karoulia et al., 2016; Wan et al., 2004; Zhang et al., 2015a). PLX7904, a modern hyperactivation paradox breaker is currently in preclinical development (Zhang et al., 2015a) while its analogue PLX8394 is in clinical trials. Here, we employ machine learning enhanced comparative analyses of MD simulation (Babbitt et al., 2018a, 2020) to identify functionally conserved loop dynamics connected to normal ATP binding. We examine how the V600E mutation breaks the natural regulatory loop dynamics of the B-Raf protein. We compare the ability of each of these four B-Raf inhibitors (see Appendix A) to mimic functionally conserved loop dynamics (i.e. creating functional dynamic mimicry (FDM) in their B-Raf targets), and we quantify the impacts of known sensitizing and drug resistant mutations on these dynamics. We also examine the V600E sensitivity of loop dynamics in two ancient phylogenetically reconstructed protein models, so as to determine what evolutionary process may have led to unusual functional vulnerability of this important signaling protein to the V600E mutation.

## Methods

### PDB structure preparation and ligand force field modification

Structures of the ATP binding domain of B-Raf bound to various Raf inhibitors were obtained from the Protein Data Bank (PDB). These were PDB ID 1uwh (wildtype B-Raf bound to sorafenib), 4rzv (vemurafenib bound to V600E mutant), 4xv1 (PLX7904 bound to V600E mutant), 4xv2 (dabrafenib bound to V600E mutant). Loop modeling and refinement were conducted on the activation loop where needed using Modeller in UCSF Chimera (Fiser et al., 2000; Pettersen et al., 2004; Sali and Blundell, 1993; Webb and Sali, 2016). Complementary mutant (V600E in 1uwh) and wildtype structures (E600V in 4rzv, 4xv1,4xv2) were computationally induced in UCSF Chimera 1.13 (Pettersen et al., 2004) by first swapping amino acids using optimal configurations in the Dunbrack rotamer library, then using 2000 steepest gradient descent steps of energy minimization to relax the regions around the amino acid replacement. To obtain ATP bound versions of wildtype B-raf and the V600E mutant, the sorafenib inhibitor was deleted from 1uwh and 1uwj, then ATP was docked using the AutoDock Vina program (Trott and Olson, 2010) after reducing both structures (adding H). ATP docking positions most closely overlapping sorafenib were chosen. Unbound forms of all the structures were obtained by deleting the ligands in the bound structure and performing energy minimization for 5000 steps. Prior to molecular dynamic simulation in Amber18, the GAFF2 force field modifications for ATP and the small molecule ligands were generated using sqm, a semiempirical and DFTB quantum chemistry program employed by the Antechamber (Ambertools18) software suite (Wang et al., 2006, 2004).

### MD simulation protocols

Large ensembles of graphic processing unit (GPU) accelerated molecular dynamic simulations were prepared and conducted using the particle mesh Ewald method employed by pmemd.cuda in Amber18 (Case et al., 2005; Salomon-Ferrer et al., 2013) via the DROIDS v3.0 interface (<u>D</u>etecting <u>R</u>elative <u>O</u>utlier <u>I</u>mpacts in <u>D</u>ynamic <u>S</u>imulation)(Babbitt et al., 2018a, 2020). Simulations were run on a Linux Mint 19 operating system mounting two Nvidia Titan Xp graphics processors. Explicitly solvated protein systems were prepared using tLeAP (Ambertools18) using the ff14SB protein force field (Maier et al., 2015) in conjunction with modified GAFF2 small molecule force field (Wang et al., 2004). Solvation was generated using the Tip3P water model (Mao and Zhang, 2012) in a 12nm octahedral water box and subsequent charge neutralization with Na+ and Cl− ions. After energy minimization, heating to 300K, and 10ns equilibration, an ensemble of 200 MD production runs each lasting 0.5 ns of time were created for both ligand bound and unbound B-Raf. Each MD production run was preceded by a single random length short spacing run selected from a range of 0 to 0.25ns to mitigate the effect of deterministic chaos (i.e. sensitively to initial conditions) in the driving ensemble differences in the MD production runs (Babbitt et al., 2018b). All MD was conducted using an Andersen thermostat (Andersen, 1980) under constant pressure. Root mean square atom fluctuations (rmsf) were calculated using the atomicfluct function in CPPTRAJ (Roe and Cheatham, 2013).

### Comparative dynamics of bound and unbound functional states

The signed symmetric Kullback-Leibler (KL) divergence (i.e. relative entropy) between the distributions of atom fluctuation (i.e. root mean square fluctuation or rmsf taken from 0.01 ns time slices of total MD simulation time) on ligand bound and unbound B-Raf were computed using DROIDS v3.0 (Babbitt et al., 2020, 2018a). This difference measure for *rmsf* has an advantage over a simple average or mean *rmsf* difference in that it captures comparative differences in the spread or deviation as well as the shape or skew of the *rmsf* distributions in addition to mean. The KL divergence between the vibrational states of two homologous atoms from two simulations, each representing a functional state of the protein, can be given by

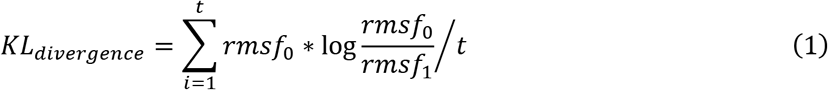

where *rmsf* represents the average root mean square deviation of a given atom over time t and 0 and 1 represent the two ensembles that are compared (i.e. in this case, the unbound and ATP bound protein states respectively). More specifically *rmsf* is a directionless root mean square fluctuation sampled over an ensemble of MD runs with similar time slice intervals. Because mutational events in evolution replace entire residues, this calculation is more useful if applied to resolution of single amino acids rather than single atoms. Because only the 4 protein backbone atoms (N, Cα, C and O) are homologous between residues, the following equation was applied. Because the remaining atoms all attach to this backbone, *rmsf* still indirectly samples the dynamic effect of amino acid sidechain replacement.

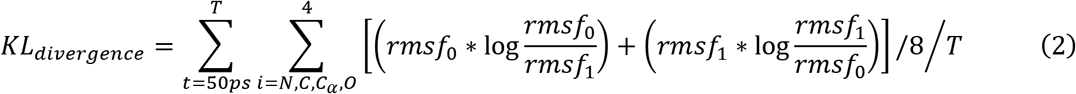

Finally, this KL divergence is made symmetric by averaging it with the KL divergence obtained when the *rmsf*_*0*_ is interchanged with *rmsf*_1_. Then the KL divergence is signed + or − depending upon whether the average atom motion (mean *rmsf*) is amplified or dampened when compared. This signed symmetric KL divergence was color mapped to the protein backbone with individual amino acid resolution to the bound structures using a temperature scale (i.e. where red is amplified fluctuation and blue is dampened fluctuation). More details of *rmsf* calculation can be found in our DROIDS v3.0 software publication (Babbitt et al., 2020). Here, the reference state of the protein is unbound while the query state is bound. Therefore, this pairwise comparison is used to represent the functional impact of ligand binding on the protein’s normal unbound motion, where it is expected that ligand contacts would typically dampen the fluctuation of atoms around the ATP binding pocket to some degree. As ATP binding acts to destabilize the activation loop and P-loop regions of B-Raf, this analysis will allow quantification of dynamic impacts that amplify loop motions as well.

### Machine learning performance and detection of conserved dynamic function

We applied machine learning classification to the dynamics of each amino acid backbone segment to differentiate functionally conserved ATP binding dynamics from unbound dynamics in repeated MD validation runs on the model of wildtype B-Raf. We then compare these results to a similar analysis of ATP binding in V600E mutants to ascertain how this mutation alters functionally conserved dynamics in the B-Raf protein system. The detection of functionally conserved protein dynamics in wildtype B-Raf followed the machine learning-based method maxDemon v1.0 outlined in our previous software paper (Babbitt et al., 2020). In summary, a stacked machine learning model implementing seven classification methods (K nearest neighbors, naïve Bayes, linear/quadratic discriminant analysis, support vector machine, random forest and adaptive boosting) was individually trained upon each amino acid’s *rmsf* data that was pre-classified to the ligand bound and the unbound dynamic states (Figure 2A). These learners were deployed upon the identical amino acids within two new 5ns MD simulations of the ligand bound structure and classifications of atom fluctuation (i.e. rmsf calculated for 500 × 0.01 ns time slices) where 0 = unbound and 1 = bound states were averaged to obtain a learning performance value for each amino acid. Average learning performance profiles (i.e. accuracy and precision) are generated along the protein from N to C terminus (Figure 2B top). Therefore, the machine learning performance at a given residue position L is given by the frequency of classification (c) where c_i_ = 0 or c_i_ = 1 taken over *t* time slices on MD simulation.

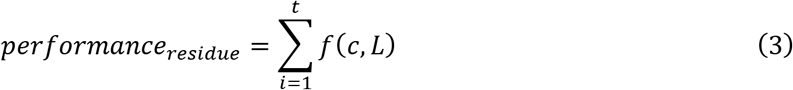

**Figure 2.**
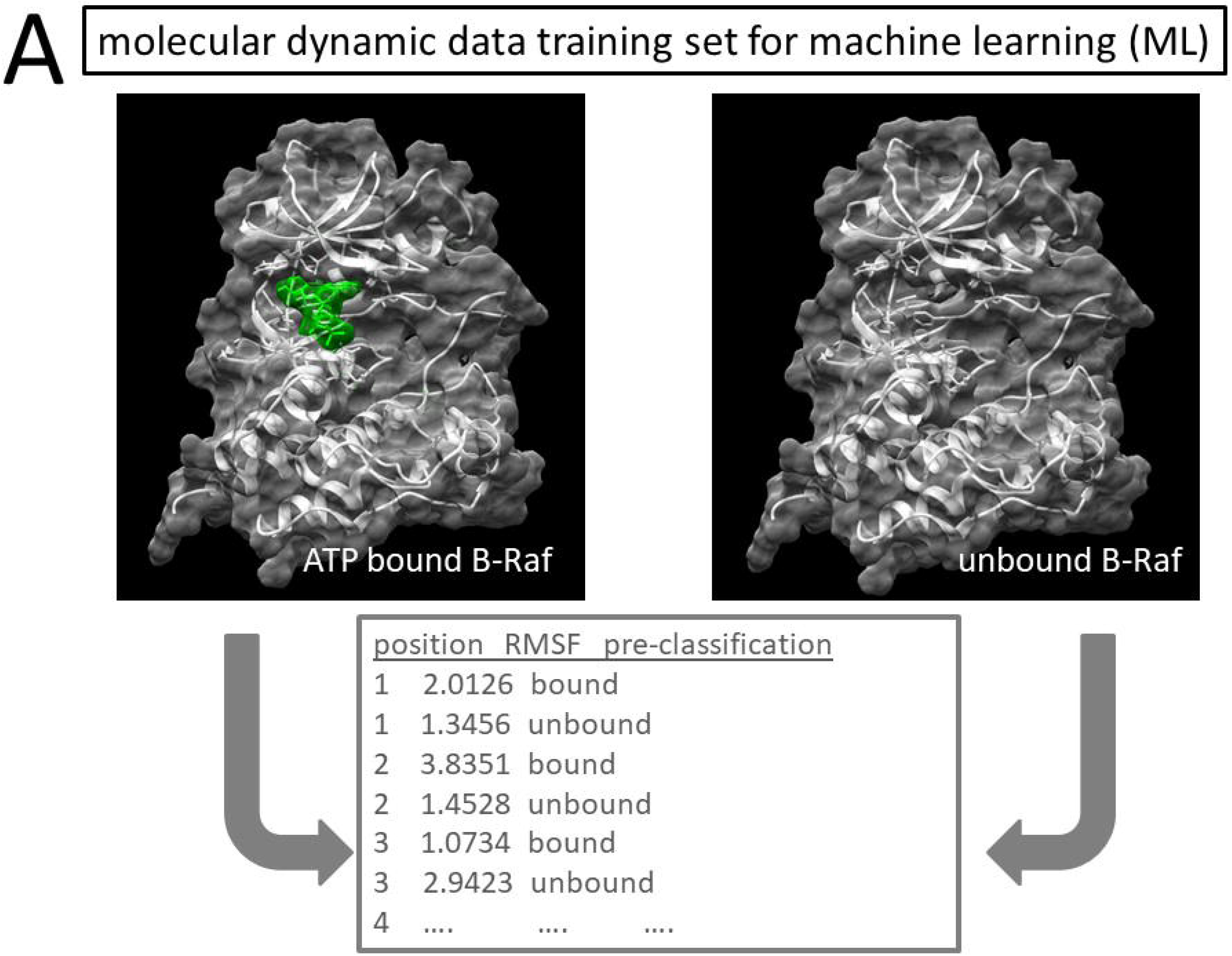
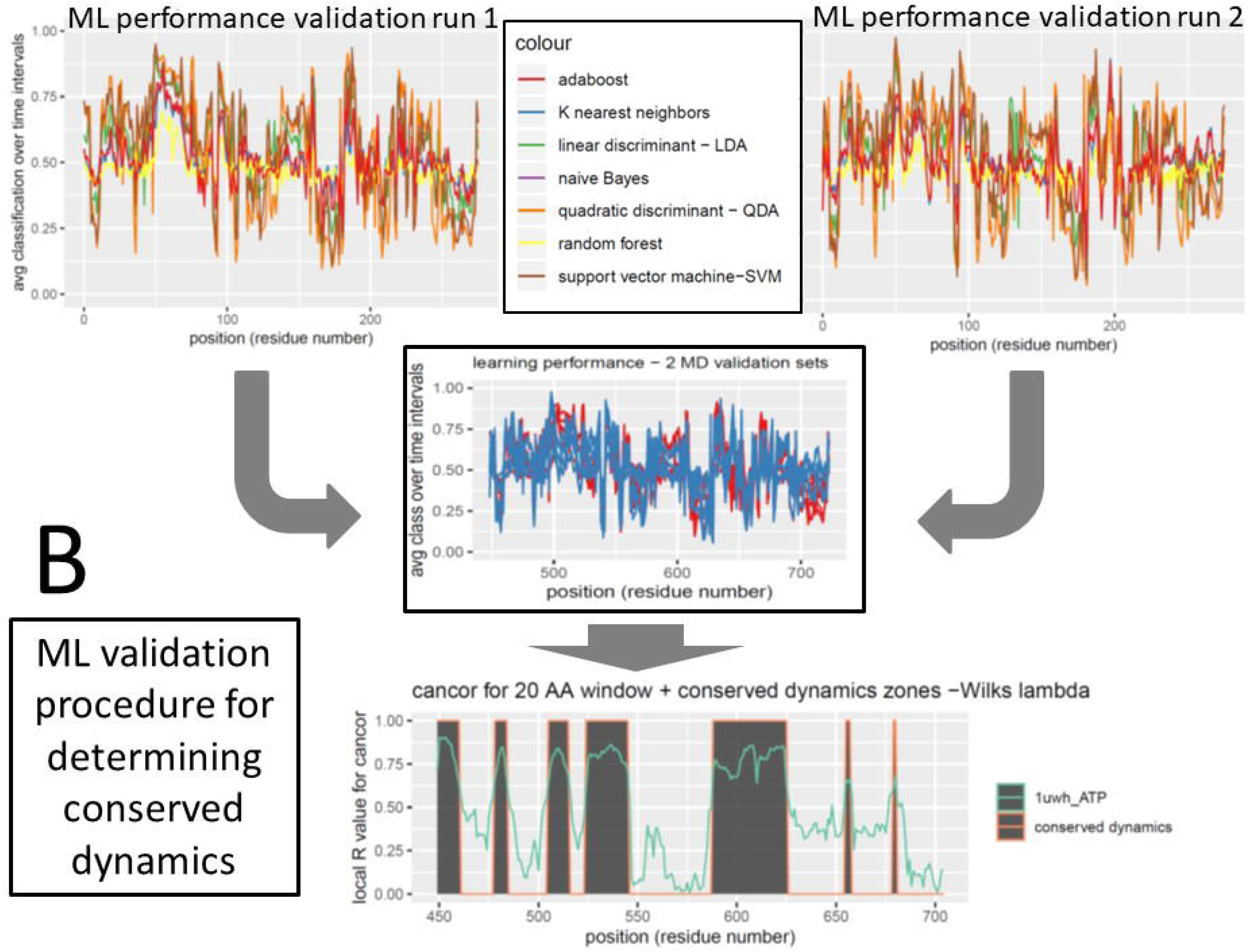
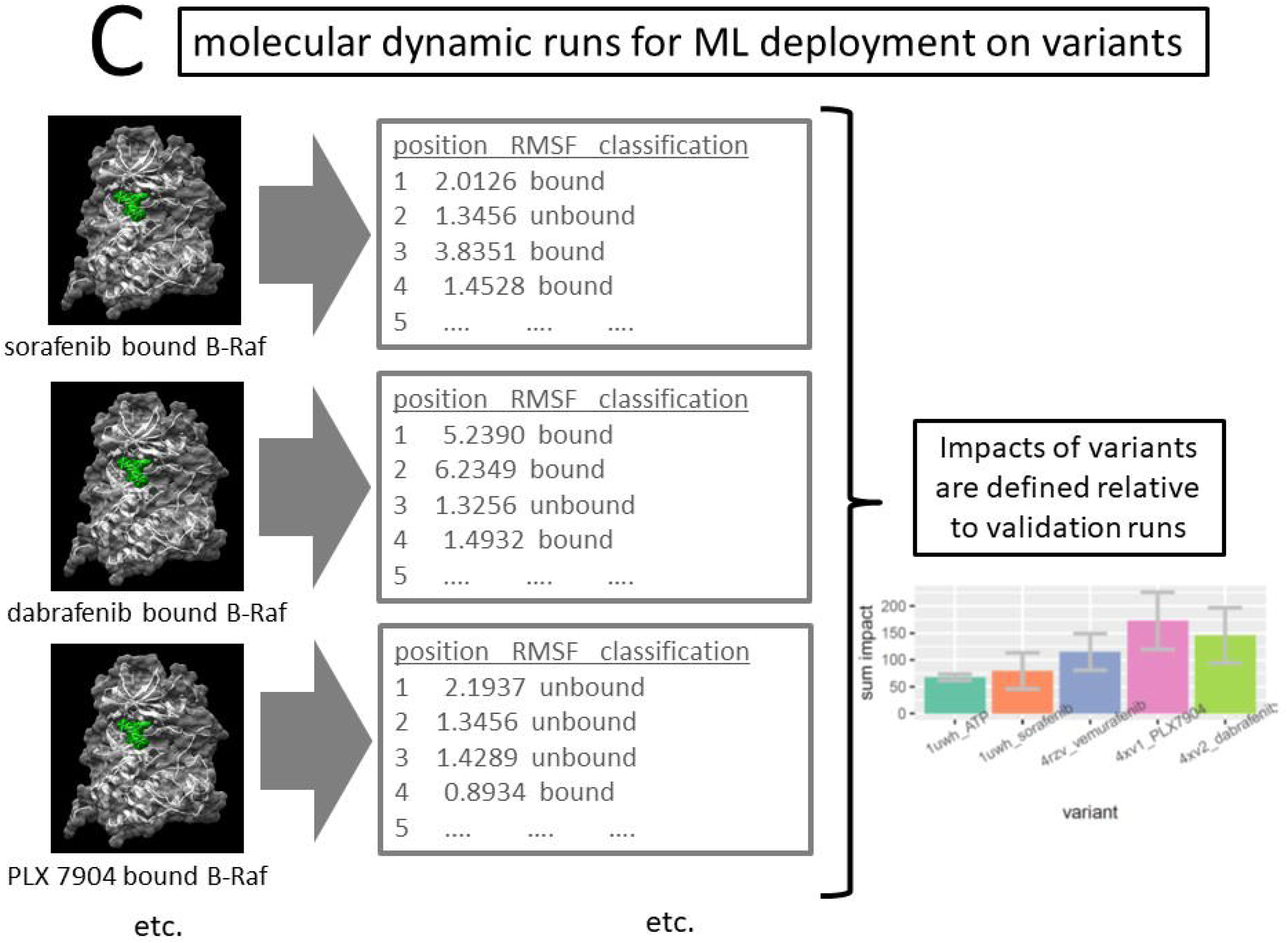
Overview of machine learning method for detecting functionally conserved B-Raf dynamics and drug class and genetic variant impacts on the conserved dynamics. (A) Pre-classified machine learning training sets are generated by large ensembles of molecular dynamic simulation of wildtype B-Raf in its two functional states; ATP bound and unbound. (B) A stacked machine learning model to identify functionally conserved molecular dynamics over two independent but identically prepared molecular dynamic validation runs on ATP bound wild type B-Raf. Position dependent machine learning performance is evaluated over time and plotted for each for each run machine learning method. The machine learning methods are listed in the center legend. Note that performance values close to 0.5 indicate random classification (i.e. no preference for either 0 or 1 classifications over time). A canonical correlation analysis of machine learning performance between the two validation runs indicates where the local classifications of dynamic functional states (i.e. representing ATP bound or unbound dynamics) are consistently repeated or ‘functionally conserved’. (C) Impacts on dynamics are determined by deploying the learner on molecular dynamic simulations of variants and then calculating the relative entropy between the dynamics of each variant and the validation runs (from B).

Note that a value of 0.5 at a given position would indicate that the learning model could not differentiate between bound and unbound dynamics at that residue. Performance profiles are generated within a 20 residue sliding window for each of the seven machine learning methods to create a composite line plot for a given MD validation or MD variant run. Local significant canonical correlations in learning performance profiles between the two 5ns MD validation runs on the ATP ligand bound structure indicate where any position-dependent molecular dynamics that is functionally conserved with regards to the ATP binding interaction has occurred(Figure 2B middle). The regions of significant functional dynamics on the B-Raf structure were called where the p-value derived from Wilk’s lambda from the local canonical correlation analysis was less than 0.01 (Figure 2B bottom). These regions are also presented with local r value from the CCA as well. Thus,

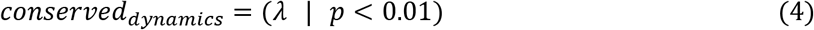

The central philosophy here is that if any atom motion is specific to a given functional state (i.e. in this case, B-Raf protein bound to ATP), then after training, the machine learners will be able to successfully classify this motion whenever it is observed in subsequent identical validation MD runs. When the learners are able to repeatedly identify any ATP-bound functional state dynamics that locally associate with any given position on the protein, this will create strongly correlated profiles in learning performance as a 20 residue site sliding window sweeps across functional regions. As our machine learning model is a stacked model, incorporating seven learning algorithms, it is more robust to artifacts related to any single method, and the resulting correlation is therefore canonical as well. We can label the dynamics as ‘functionally conserved’ if the machine learning model can consistently identify the functional dynamic states upon which it trained across multiple and independent validation runs that are of adequate length (i.e. many nanoseconds). Two or more identically prepared validation runs are used to examine whether statistically significant canonical correlations (i.e. conserved dynamics) are locally identifiable by a significant local Wilk’s lambda value for a local canonical correlation coefficient.

### Method of genetic and drug variant impact assessment

The local impacts of mutations and drug class variants were obtained by first deriving the canonical correlation of the learner performance profile of a given variant to the wildtype B-Raf in its normal functionally bound state (= CC_variant_). This also uses a 20 residue sliding window. The impact metric is generated by comparing canonical correlation of the variant (CC_variant_) to the self-similar dynamic correlation (CC_self_) defining functionally conserved dynamics above, using the common definition of relative entropy below.

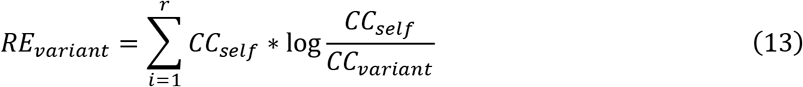

Effectively, this impact metric captures the entropy difference in conserved dynamics between the wildtype ligand bound state (i.e. ATP bound) and the variant states of binding created by genetic mutation or different drug class ligand molecules. This occurs on the same scale as that used to differentiate bound from unbound states during the earlier training of the stacked learning model. More details regarding our methodological approach to identifying functionally conserved dynamics and subsequent impacts of genetic and drug class variants can be obtained in our prior publication (Babbitt et al., 2020). The error bars shown in relation to drug class and genetic variants impacts across athe protein are derived by 1000 bootstrapped samples of the local machine learning classifier outputs (i.e. 0 or 1) taken at all amino acid residue sites.

The relative impacts of four B-Raf inhibitors, sorafenib, dabrafenib, vemurafenib and PLX7904 were assessed after initially training the stacked machine learning model on ATP bound and unbound B-Raf (Figure 2C) within both the wildtype and the V600E mutant genetic background. Several drug affecting genetic mutants were also examined after training the learning model on normal functional binding interaction of each drug in the wildtype genetic background. These drug resistant and drug sensitizing mutations were identified using the Variant Interpretation for Cancer Consortium (VICC) Meta-Knowledgebase (Wagner et al., 2018). Variants were selected that contained literature support for the functional impact on at least one of the four B-Raf inhibitors (Supplemental Table 1).

### Phylogenetic analysis of B-Raf sensitivity to V600E

A set of 27 BRAF ortholog sequences were obtained using the National Center for Biotechnology Information (NCBI) ortholog finder tool. These sequences were aligned using MEGA 7.0 (Kumar et al., 2016) Clustal multiple sequence alignment tool and then manually trimmed to contain only the ATP binding domain. Phylogenetic model selection was also conducted in MEGA to identify the most likely (best fitting) evolutionary substitution model. This was identified via both Bayesian and Akaike Information Criterion to be the Kimura 2-parameter model with extra terms for both gamma correction for multiple same site substitution and correction for invariants sites (i.e. K2+G+I). A minimum evolution tree and a maximum likelihood tree bootstrapped 500 times confirmed the same tree topology with high confidence. Amino acid replacements distributed on this topology identified three protein variants, one each for tetrapods, ray-finned fish and insects. Amino acids differences from tetrapods identified 9 amino acid replacements since a common ancestor with ray-finned fish and 95 replacements since a common ancestor with insects. Structural models for the two ancestral protein variants were manually generated in UCSF Chimera using swapaa commands to find non-clashing Dunbrack library rotamer configurations. These models were then subsequently relaxed with 2000 steps of energy minimization via steepest gradient descent. A V600E variant was subsequently created from the wildtype ray-finned fish model and an A600E variant was created from the wildtype insect model. DROIDS software was used to train upon ATP binding interaction in the wildtype tetrapod model (comparing ATP bound and unbound dynamics as described above). DROIDS 3.0 and maxDemon 1.0 machine learning analysis (described above) was then used to identify functionally conserved dynamics in tetrapod B-Raf and subsequently compare mutational impacts on B-Raf dynamics for both the ancestral wildtype and mutant backgrounds in ray-finned fish and insects.

The author’s software can be accessed at the software landing website and GitHub repository link

https://people.rit.edu/gabsbi/

https://github.com/gbabbitt/DROIDS-3.0-comparative-protein-dynamics

All data for the analyses and Figures are available at Zenodo (search title of paper), an open access data repository hosted at CERN.

## Results

### Shifts in B-Raf loop dynamics upon ATP binding in wildtype and V600E mutant background

The comparative analysis of wildtype B-Raf dynamics (rmsf) in both its ATP-bound and unbound form revealed upon binding, a significant dampening of atom fluctuation in the ATP binding pocket associated with highly pronounced increase in fluctuation in the region of the activation loop as well as upstream toward the C terminal approximately affecting residues 599-625 (Figure 3A, 2C, Supplemental Figure 1A-C). Note: this shift in dynamics was indicated by decreased rmsf or negative dFLUX defined by the KL divergence in rmsf. Shifts in dynamics were significant according to multiple test corrected Kolmogorov-Smirnov tests at four major regions (Supplementary Figure 1C) including the binding pocket, the catalytic + activation loops and at two regions centered at residue 550 and 660. When color mapped to the ATP bound structure, these dynamics exhibit a large physical separation between the activation loop and P-loop (i.e. open active conformation for the protein) (Figure 3A).

**Figure 3.**
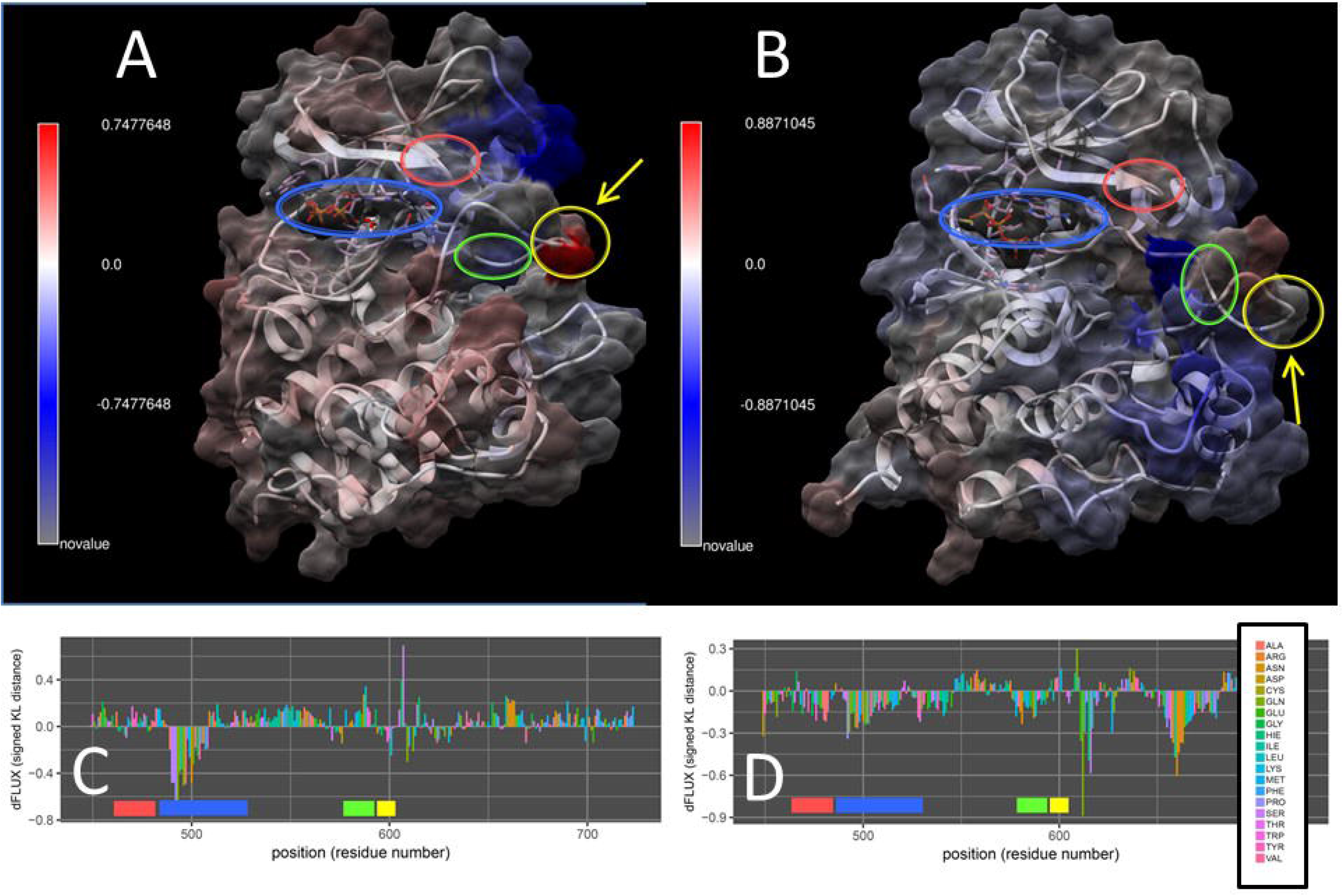
Effect of V600E mutation on rapid dynamics of ATP binding interaction in human BRAF kinase domain. Signed symmetric Kullback-Leibler (KL) divergences (i.e. relative entropy) in ATP-bound and unbound empirical distributions of local root mean square fluctuation (i.e. dFLUX) derived from large ensembles of nanosecond scale molecular dynamics simulations (300×0.5ns) are shown color mapped on the (A) ATP-docked wild type BRAF (ATP docked to PDB ID 1uwh receptor) and the (B) V600E mutant BRAF structure (ATP docked to PDB ID 1uwj receptor) with red indicating amplified dynamics upon ATP binding and blue indicating stabilized dynamics upon ATP binding. Plots of signed symmetric KL divergence for (C) wild type BRAF and (D) V600E mutant BRAF are also shown. Positive KL divergence indicates amplified dynamics upon ATP binding while negative values indicates stabilization upon binding. Colored circled regions (A-B) and bars (C-D) indicate the P-loop (red), ATP binding pocket (blue), catalytic loop (green) and a highly flexible activation loop (yellow). Yellow arrows also indicate location of the start of the activation loop and highlight a highly pronounced increase in flexibility of this region during ATP binding in wild type BRAF that contrasts a dysfunctional stabilization of this region and areas downstream in the V600E mutant.

Comparative dynamics analysis of same ATP binding interaction in the presence of the V600E mutation exhibited clear and striking differences in dynamic shift (i.e. KL divergence) compared to the wildtype background. ATP binding in this mutant not only stabilized the regions around the P-loop and ATP binding pocket, but also massively and significantly stabilized the dynamics of the region of the catalytic + activation loop and a large region from residue 650-680 (Figure 3B, 2D, Supplemental Figure 2 A-C) serving to apparently freeze the activation loop into an open configuration characteristic of the V600E mutant (Figure 3B).

### Functionally conserved B-Raf loop dynamics upon ATP binding in wildtype and V600E mutant background

Our machine learning model trained to recognize functional binding interaction dynamics in ATP bound wildtype B-Raf revealed clear non-random patterns of learning performance at specific residue locations on the protein that were highly correlated across the two MD validation runs. Significant and strong canonical correlations in learning performance on the MD validation runs indicated conserved binding dynamics across the regions connecting the ATP binding pocket to both the tip of the P-loop and a large region surrounding the catalytic and activation loops (Figure 4A and 4C). High R values in these regions indicate that the learning algorithms identified dynamics characteristic of B-Raf’s interaction with ATP over 65% of the total time of simulation, in spite of the thermal fluctuation that is always present. The plots of local learning performance of each algorithm in each of the two validation runs also indicates that characteristic functional ATP binding dynamics in B-Raf corresponded well with the known functional regions of the protein as well as the C terminal region approximately from residue 625-675.

**Figure 4.**
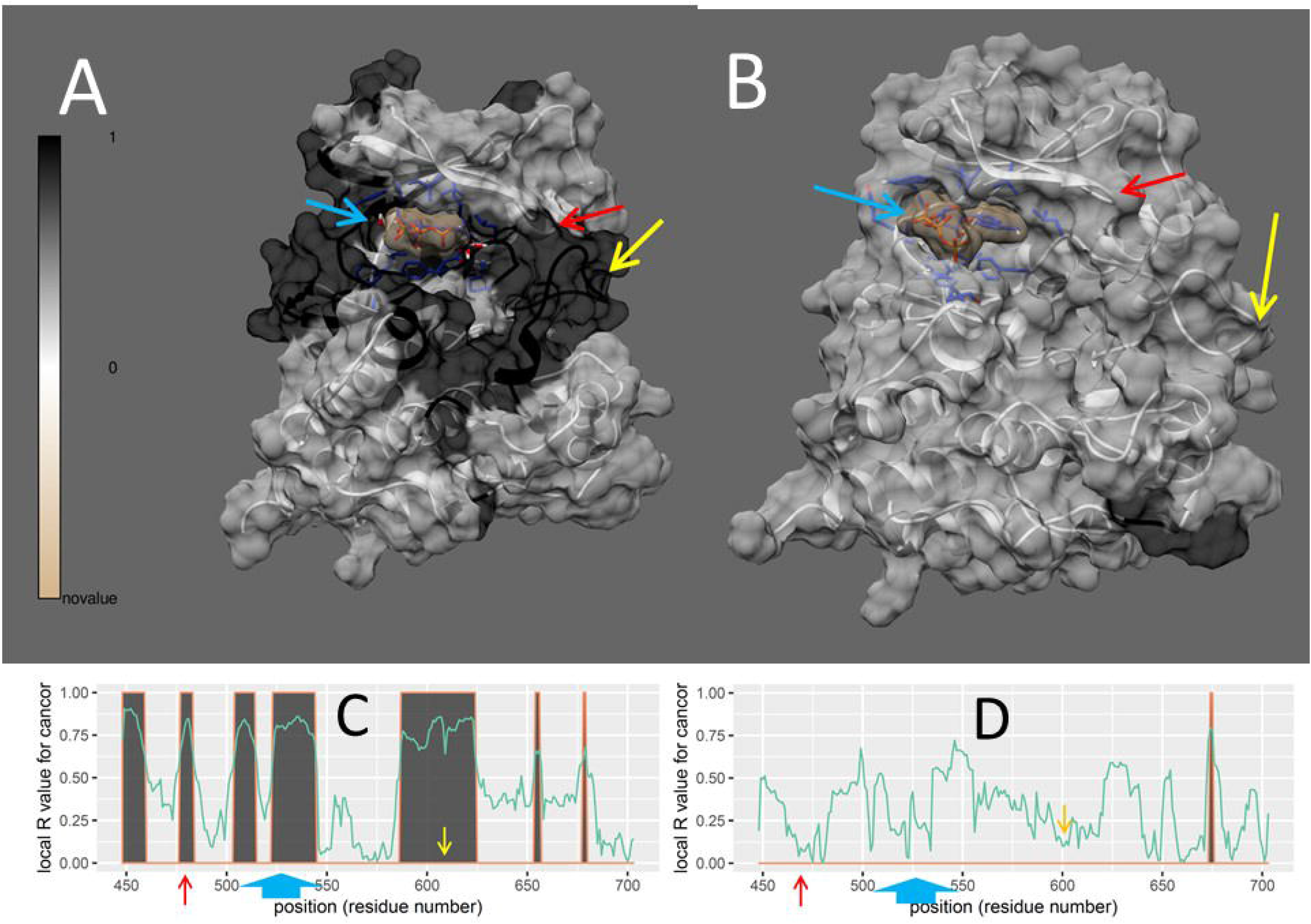
Effect of V600E mutation on functionally conserved dynamics of ATP binding interaction in human BRAF kinase domain. (A) In wild type BRAF, the machine learning model identifies functionally conserved (i.e. correlated) molecular dynamics that clearly connect the ATP binding pocket (blue arrow; ATP structure is tan) to the activation loop (yellow arrow), p loop (red arrow) and surrounding structures. Significantly conserved local dynamic regions determined via Wilk’s lambda for the canonical correlation analysis are shown in dark gray. A 20 residue sliding window was used to the statistic. (B) In V600E mutant BRAF, the functionally conserved dynamics of ATP binding are almost completely absent. The local R values and significance are also plotted for (C) wild type BRAF and (D) V600E mutant BRAF. Again dark gray indicates regions of significant conservation.

When deployed upon V600E mutant model simulations, the multi-machine learning algorithm generally failed to find conserved ATP binding dynamics (Figure 4B and 4D) typical of normal wildtype B-Raf. In this malfunctional genetic background, low r values and lack of significance in the canonical correlation analysis of learning profiles indicate that our machine learning model fails to identify any functionally conserved dynamics of ATP binding, except in a very small part of the region (650-680) that freezes to hold the activation loop (Figure 4D). The local machine learning performance was generally poorer across the whole domain (i.e. closer to 0.5) indicating that functional dynamics played a far less important functional role in the V600E mutant background than in wildtype (Supplemental Figure 3). Taken altogether this result is likely indicative of how B-Raf’s very low tolerance to V600E destroys the functional dynamics of ATP binding in B-Raf.

### Effects of Raf inhibitors on conserved B-Raf loop dynamics in wildtype and V600E mutant background

Because we theorized that the hyperactivation of cancer in wildtype cells by certain B-Raf inhibitors could be due to their potential to mimic the natural agonist of MAPK (i.e. ATP), we deployed our multi-machine learning model, already trained upon dynamic differences caused by functional ATP interaction in wildtype B-Raf, upon models of wildtype B-Raf bound to four specific B-Raf inhibitors. These inhibitors included sorafenib, a potent inducer of hyperactivation and PLX 7904, a hyperactivation paradox breaker, currently in pre-clinical stages of research. We also include two modern B-Raf inhibitors, currently used in combination in clinical settings. We find sorafenib closely mimics ATP both in its pattern of functionally conserved dynamics (Figure 5A) and in the magnitude of its dynamic impacts on conserved dynamics of the total protein (Figure 5B). Here, the more modern inhibitors, vemurafenib, dabrafenib and PLX7904 show fewer regions with significantly conserved dynamics and higher associated impacts on conserved dynamics. The highest dynamic impact was in PLX7904, a potent ‘breaker’ of the ‘paradox of hyperactivation’, suggestive that unlike sorafenib, its binding to wildtype B-Raf may even serve to render B-Raf unrecognizable to the MAPK pathway. Plots of these conserved dynamic regions and their representative local r values are shown in Supplemental Figure 4. Local plots of relative impacts of each inhibitor in both wildtype and V600E mutant background are shown in Supplemental Figure 5. Comparative dynamic analysis of *rmsf* for B-Raf bound to each of the four inhibitors within wildtype backgrounds (Supplemental Figure 6) also indicates that sorafenib most closely mimics ATP via its interaction with the binding pocket and subsequent amplifications of dynamics of the activation loop (i.e. see similarity between Figure 2B and Supplemental Figure 6A). While the modern inhibitors have KL divergence profiles somewhat similar to that of ATP in wildtype backgrounds, they all differ in key respects. Vemurafenib and dabrafenib also seem to affect P loop dynamics in ways that are very different than ATP (position 460-468 in Supplemental Figure 6B and 6C), while vemurafenib and PLX7904 exhibits more global disruption and dampening of dynamics across the whole B-Raf binding domain (indicated by widespread negative KL divergence in Supplemental Figure 6B and 6D). We note that PLX7904, as the paradox breaking inhibitor, shows substantially less activity near the V600 region and activating segment in general, as it was constructed to disrupt overall B-Raf structure and affect dimerization, particularly targeting the L505 position, in order to affect the helix there. In our study, we find that the region near 490-505 appear to be selectively affected by disruptions to dynamics, suggesting a dynamic mechanism for its previously observed hyperactivation paradox breaking behavior.

**Figure 5.**
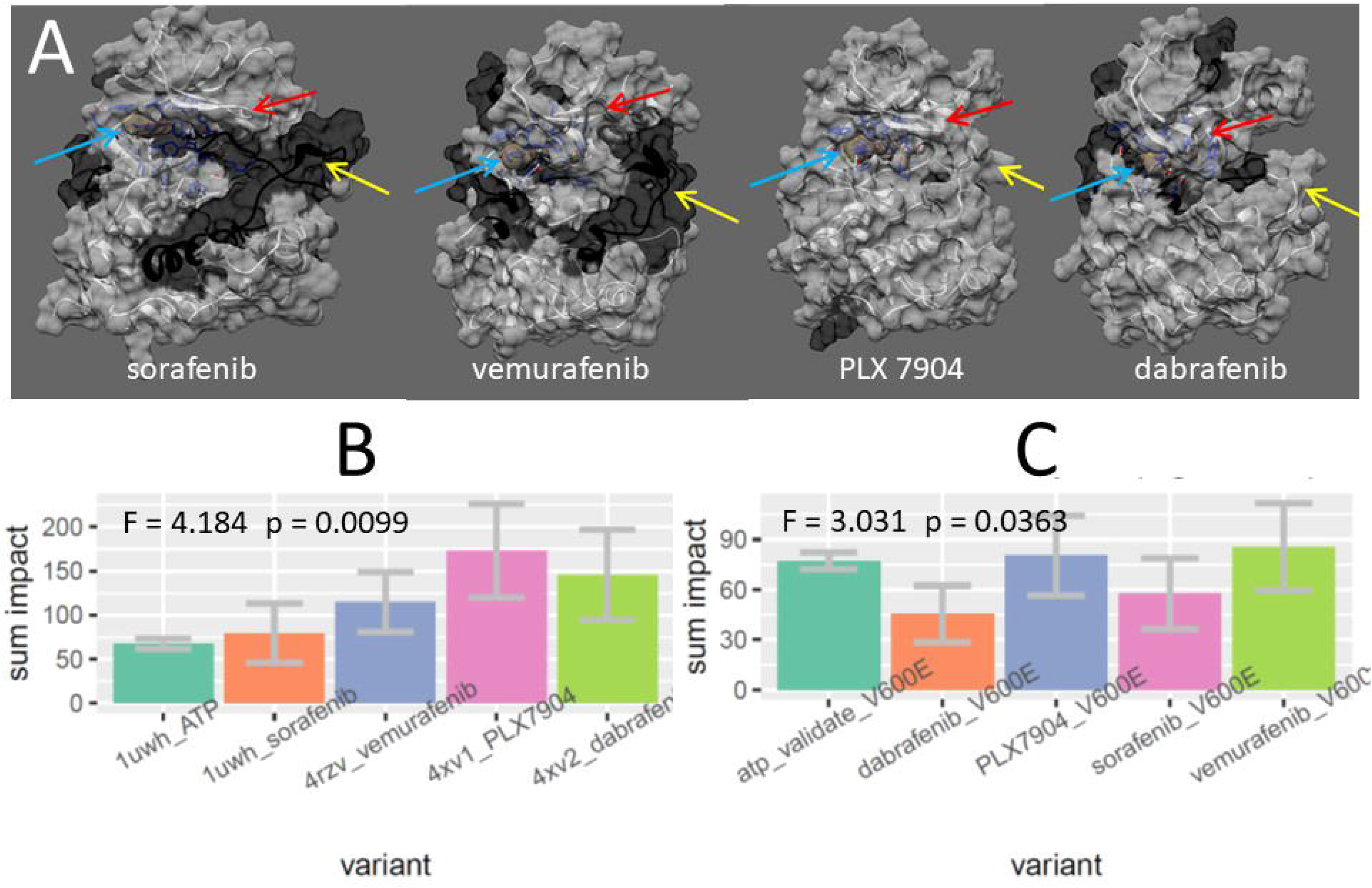
Comparisons of the effects of four BRAF inhibitors on functionally conserved dynamics of ATP binding interaction in human BRAF kinase domain. (A) Significantly functionally conserved dynamic regions are shown in dark gray for each of four BRAF inhibitors. The first generation inhibitor sorafenib seems to closely mimic ATP in its effect on wild type BRAF dynamics, while the hyperactivation paradox breaker, PLX7904 seems to mimic functional dynamics of ATP binding the least. Arrows indicate the ATP binding pocket (blue arrow; ATP structure is tan), the activation loop (yellow arrow),and the p loop (red arrow). (B) the sum of relative impacts on conserved dynamics across the wild type BRAF protein also indicates that sorafenib is most similar to ATP in its effect on wild type dynamics, while PLX7904 is least similar, with much more global disruption of conserved dynamics. (C) The same analysis conducted on the V600E mutants indicates that all four inhibitors are similar to ATP their impact on dynamics. Note that V600E will likely have eliminated conserved dynamics as in Figure 5B. Plots of the canonical correlation analysis are shown in supplemental Figure C. Mutual information matrices are shown in supplemental Figure D.

As the V600E mutant does not exhibit high levels of functionally conserved dynamics regarding ATP (Figure 4B), we would not expect B-Raf inhibitors to exhibit high levels of disruption to conserved dynamics upon binding in the V600E mutant. This is confirmed by the observation that the impacts of the four B-Raf inhibitors on conserved dynamics were roughly equivalent to that of ATP in the mutant background (Figure 5C). Comparative plots of KL divergence in *rmsf* of the four B-Raf inhibitors in the V600E mutants (Supplemental Figure 7) indicated that sorafenib globally destabilized B-Raf upon interaction with the ATP binding pocket (Supplemental Figure 7A). This effect was larger toward the C terminal regions of the protein. In the mutant background, Vemurafenib appeared to mimic the functional effect of ATP binding similar to wildtype, but too a much larger degree, confirming a potential indication of its well-known selectivity to mutant B-Raf (Supplemental Figure 7B). Upon its interaction with the binding pocket, Dabrafenib induced strong stabilization in the area of the catalytic and activation loops, locking B-Raf into its open confirmation and perhaps enhancing its well-known effect as an ATP competitor (Supplemental Figure 7C). Alternatively, upon binding the mutant, PLX7904 exhibited large destabilizations of the P-loop, ATP binding pocket and the regions extending from the activation loop and the region of residue 650-680 that freezes the open confirmation in the absence of the drug (Supplemental Figure 7D).

In summary, the hyperactivation-breaking compound, PLX7904, while also causing similar dynamics shifts to the activation loop, fails to produce hardly any dynamic changes that are identified as functional according to our machine learning classification method. Thus, it appears that while PLX7904 is a potent competitive binder to mutant B-Raf, it also may have the benefit of potentially disrupting any characteristic dynamics related to ATP binding that could allow stable interactions with other components of the MAPK pathway. Our analysis of mutational impacts on functional dynamics also support this, and suggest that changes in PLX7904 efficacy caused by mutation are likely independent of the normal evolutionary conserved function of B-Raf. Analysis of vemurafenib shows some similarity to sorafenib, suggesting that it is also prone to mimic the functional dynamics of ATP binding but with softening of the P-loop as well. Similar analysis of dabrafenib indicates a hardening of P-loop dynamics and a pattern of less coordinated impact on conserved dynamics that fails to connect the ATP binding pocket to the activation loop, supporting our hypothesis that more modern Raf inhibitors are less like ATP in their overall effect on functional shifts in B-Raf dynamics during binding.

### Comparison of impacts of drug sensitivity and resistant mutations on B-Raf loop dynamics

To further investigate the effect of both genetic and drug class variation on functional B-raf loop dynamics, we compared the dynamic impacts on B-Raf for a variety of drug sensitizing and drug resistance linked mutations for each of the four Raf inhibitors, sorafenib, vemurafenib, dabrafenib, and PLX7904 (Supplemental Table 1, Figure 6). The V600E mutation which sensitizes sofafenib targeting to B-Raf, was found to more greatly impact loop dynamics when compared to validation runs with no and corroborating its large impact on functional dynamics in general. The three remaining drug resistant mutations (G464V, G469E, G469V) show less impact, but are collectively larger than validation, suggesting that mutations disrupting dynamics of the P loop are key in creating drug resistance to sorafenib (Figure 6A). It should be noted that our bootstrapping did not find these differences to be significant. In the PLX 7904 hyperactivation breaker, it is noted that while not many drug affecting mutations have been discovered, they are mostly congruent with the mutationally intolerant region (V600) (Figure 6B). In more modern inhibitors, a survey of 18 drug impacting mutations does not show any clear trend except that mutational impacts are significantly largest in regions affecting either the P-loop or the activation loop (Figure 6C and 6D). Mutations affecting dabrafenib function also appear to be largely confined to P-Loop and activation loop, with the exception of L514V, a site whose side-chain actively engages ATP during normal B-Raf function.

**Figure 6.**
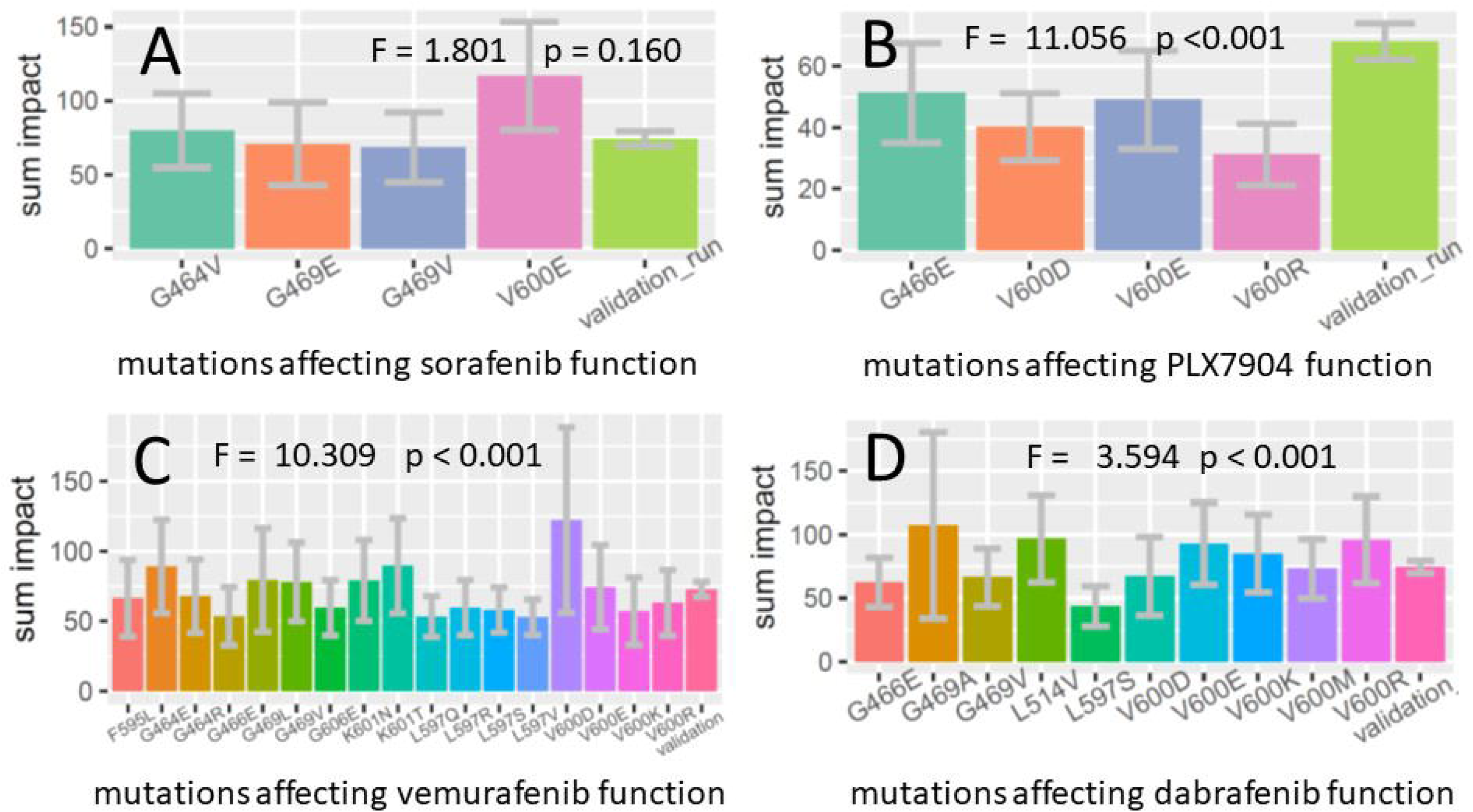
Relative impacts of drug sensitizing and drug resistant mutations on functionally conserved binding dynamics in human BRAF kinase domain. (A) The impacts of four known drug sensitizing mutations for sorafenib are compared (V600E, G469V, G469E, G464V). (B)The impacts of three known drug sensitizing mutations (V600E, V600R, V600D) and one drug resistant mutation (G466E) in PLX7904 are also compared. (C) The impacts of 18 known drug sensitizing and resistant mutations in vemurafenib 10 known drug sensitizing and resistant mutations in dabrafenib and are also compared. See Table 1 for classifications.

**Table 1.** Genetic variants affecting drug activity of four Raf inhibitors (sorafenib, vemurafenib, dabrafenib, PLX7904).

| drug | disease | mutation | response | source |
| --- | --- | --- | --- | --- |
| sorafenib | breast cancer | G464V | sensitivity | http://www.ncbi.nlm.nih.gov/pubmed/18519791 |
| sorafenib | cutaneous melanoma (disease) | G469E | sensitivity | http://www.ncbi.nlm.nih.gov/pubmed/18794803 |
| sorafenib | non-small cell lung carcinoma (disease) | G469V | sensitivity | http://www.ncbi.nlm.nih.gov/pubmed/27388325 |
| sorafenib | thyroid cancer | V600E | sensitivity | http://www.ncbi.nlm.nih.gov/pubmed/18682506 |
| PLX7904 | melanoma (disease) | G466E | resistance | http://www.ncbi.nlm.nih.gov/pubmed/27523909 |
| PLX7904 | melanoma (disease) | V600D | sensitivity | http://www.ncbi.nlm.nih.gov/pubmed/27523909 |
| PLX7904 | melanoma (disease) | V600E | sensitivity | http://www.ncbi.nlm.nih.gov/pubmed/26466569 |
| PLX7904 | melanoma (disease) | V600R | sensitivity | http://www.ncbi.nlm.nih.gov/pubmed/27523909 |
| vemurafenib | cancer | G464E | resistant | http://www.ncbi.nlm.nih.gov/pubmed/29320312 |
| vemurafenib | cancer | G464R | resistant | http://www.ncbi.nlm.nih.gov/pubmed/26343582 |
| vemurafenib | melanoma (disease) | G466E | resistant | http://www.ncbi.nlm.nih.gov/pubmed/27523909 |
| vemurafenib | cancer | G469L | conflicting | https://www.ncbi.nlm.nih.gov/pubmed/26200454 |
| vemurafenib | cancer | G469V | resistant | http://www.ncbi.nlm.nih.gov/pubmed/26343582 |
| vemurafenib | cancer | F595L | resistant | http://www.ncbi.nlm.nih.gov/pubmed/26582644 |
| vemurafenib | colorectal cancer | G596R | resistant | http://www.ncbi.nlm.nih.gov/pubmed/22180495 |
| vemurafenib | cancer | L597Q | resistant | http://www.ncbi.nlm.nih.gov/pubmed/26343582 |
| vemurafenib | melanoma (disease) | L597R | sensitivity | http://www.ncbi.nlm.nih.gov/pubmed/23715574 |
| vemurafenib | cancer | L597S | sensitivity | http://www.ncbi.nlm.nih.gov/pubmed/22798288 |
| vemurafenib | cancer | L597V | resistant | http://www.ncbi.nlm.nih.gov/pubmed/26343582 |
| vemurafenib | cancer | G606E | No benefit | http://www.ncbi.nlm.nih.gov/pubmed/29320312 |
| vemurafenib | cancer | K601N | resistant | http://www.ncbi.nlm.nih.gov/pubmed/26343582 |
| vemurafenib | cancer | K601T | resistant | http://www.ncbi.nlm.nih.gov/pubmed/26343582 |
| vemurafenib | melanoma (disease) | V600D | sensitivity | http://www.ncbi.nlm.nih.gov/pubmed/27523909 |
| vemurafenib | melanoma (disease) | V600E | sensitivity | http://www.ncbi.nlm.nih.gov/pubmed/27297867 |
| vemurafenib | melanoma (disease) | V600K | sensitivity | http://www.ncbi.nlm.nih.gov/pubmed/20149136 |
| vemurafenib | melanoma (disease) | V600R | sensitivity | http://www.ncbi.nlm.nih.gov/pubmed/27523909 |
| dabrafenib | melanoma (disease) | G466E | resistant | http://www.ncbi.nlm.nih.gov/pubmed/27523909 |
| dabrafenib | non-small cell lung carcinoma (disease) | G469A | sensitivity | http://www.ncbi.nlm.nih.gov/pubmed/29903896 |
| dabrafenib | breast cancer | G469V | No benefit | http://www.ncbi.nlm.nih.gov/pubmed/24885690 |
| dabrafenib | breast cancer | L514V | resistant | http://www.ncbi.nlm.nih.gov/pubmed/29880583 |
| dabrafenib | melanoma (disease) | L597S | sensitivity | http://www.ncbi.nlm.nih.gov/pubmed/29903896 |
| dabrafenib | melanoma (disease) | V600D | sensitivity | http://www.ncbi.nlm.nih.gov/pubmed/27523909 |
| dabrafenib | melanoma (disease) | V600E | sensitivity | http://www.ncbi.nlm.nih.gov/pubmed/20818844 |
| dabrafenib | melanoma (disease) | V600K | sensitivity | http://www.ncbi.nlm.nih.gov/pubmed/21343559 |
| dabrafenib | melanoma (disease) | V600M | sensitivity | http://www.ncbi.nlm.nih.gov/pubmed/23031422 |
| dabrafenib | melanoma (disease) | V600R | sensitivity | http://www.ncbi.nlm.nih.gov/pubmed/27523909 |

### Phylogenetic analysis of V600E sensitivity regarding B-Raf loop dynamics

A minimum evolution tree (Figure 7A) and maximum likelihood tree (supplemental Figure 8) was used to identify protein level variants for tetrapod (= human), ray-finned fish, and insect. The wildtype genetic backgrounds incorporating 9 amino acid replacements (fish) and 95 replacements (insect) show no elevated dynamic impacts over the validation run (wildtype tetrapod). However, V600E/A600E mutants both demonstrate elevated disruption of conserved ATP binding dynamics (Figure 7B) indicating that B-Raf sensitivity to this mutation likely has an ancient Eukaryotic origin. The very large increase in dynamic impact of V600E in ray-finned fish over that of A600E in insects also strongly indicates a subsequent increase in V600E sensitivity in early jawed vertebrate evolution as well (Figure 7B). Local mapping of these impacts (Figure 7C) indicates involvement of all three functional regions (P-loop at position 13-25, ATP binding pocket at position 60-100, and activation loop at position 153-190) in the functional evolution of this sensitivity.

**Figure 7.**
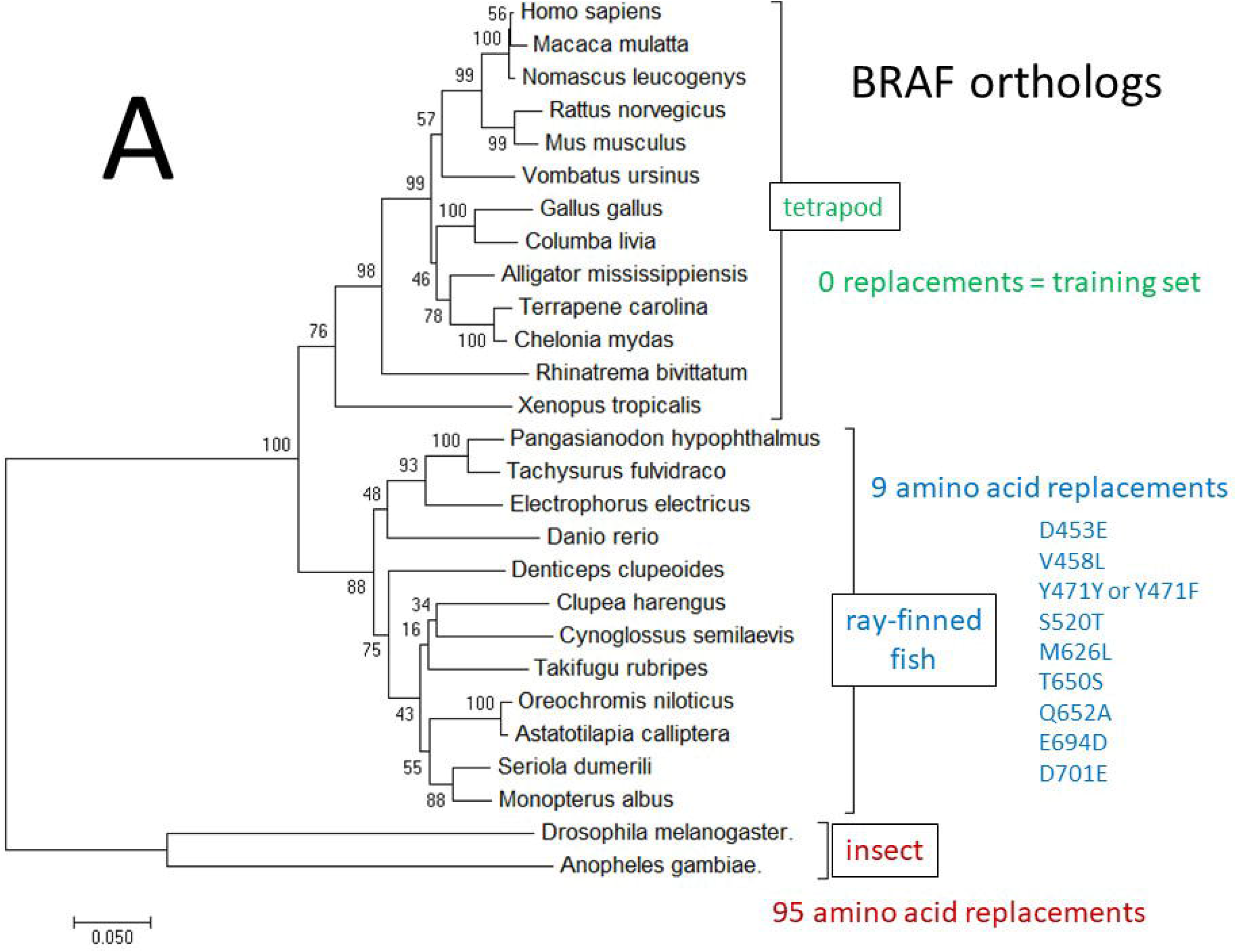
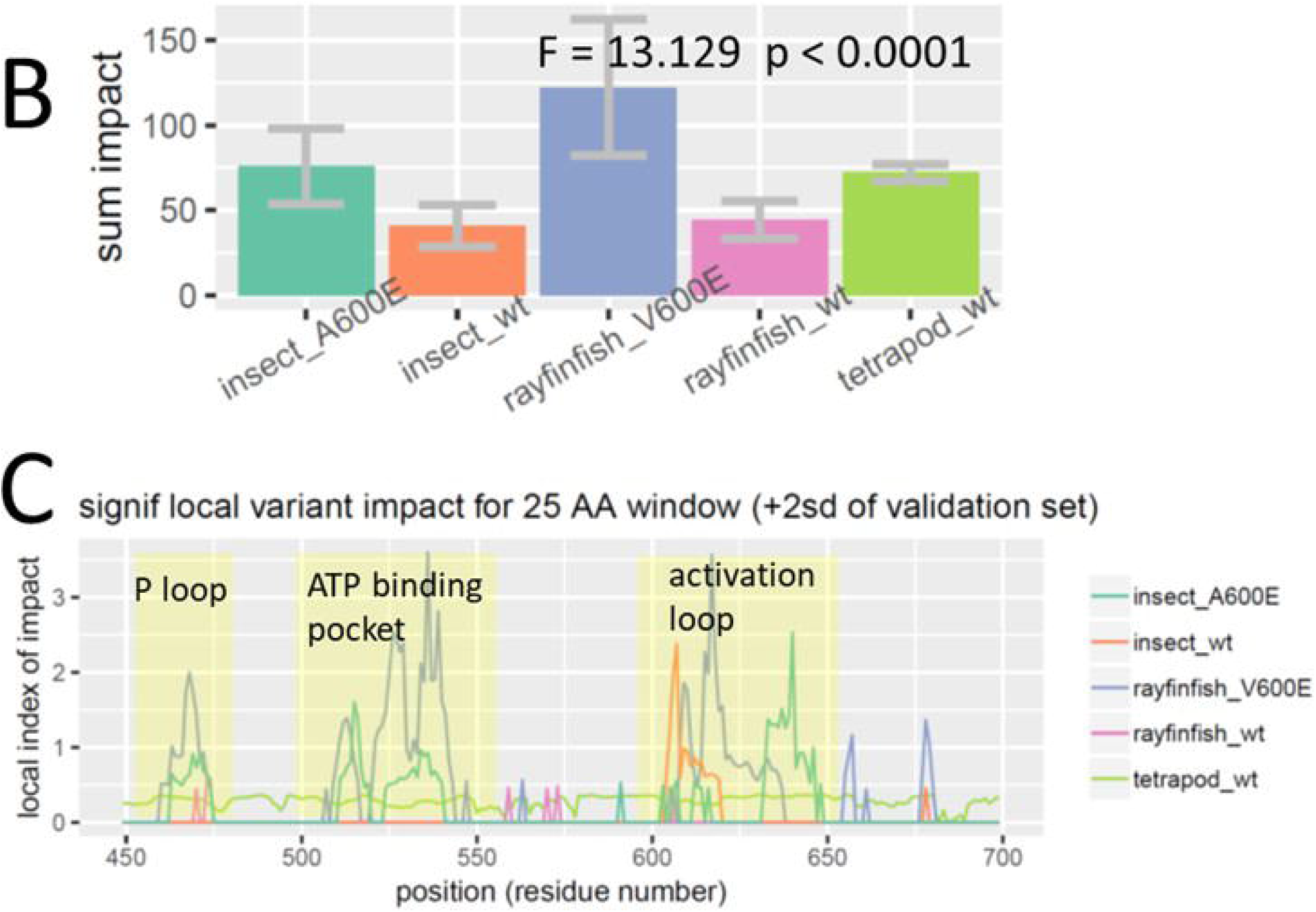
Comparative functional molecular dynamic analysis of ATP binding in wild type BRAF orthologs. Orthology was determined via phylogenetic analysis to derive a (A) minimum evolution tree bootstrapped 500 times using the Kimura 2 parameter model with Gamma correction for multiple hits and invariant sites. This best model was determined via multi-model inference (BIC, AICc). (B-C) With a goal of determining when V600E sensitivity first arose in this phylogeny, we compare the conserved region impacts of ATP binding on BRAF dynamics in wild type and V600E mutated genetic backgrounds of ray-finned fish and insect ortholog models after training on the functional ATP binding in wild type tetrapod (i.e. human) background. These impacts are shown both (B) globally and (C) locally, where regions of low mutational tolerance are observed corresponding to the P-loop, ATP binding pocket, and activation loop and adjacent downstream residues (shaded yellow).

## Discussion

Given their involvement in a variety of cellular processes, protein kinases, specifically serine/threonine kinases play a role in all of the major cancers. Many serine/threonine kinases have altered expression in tumors (Capra et al., 2006). Given that this class of enzyme is a proven putative and attractive target for the development of cancer therapies, our study of the function and evolution of the molecular dynamics of the serine/threonine kinase B-Raf has the potential to guide future studies and experiments aimed at elucidating efficacy and rationale design of therapeutics. Through our novel high performance computing approach to machine learning applied to comparative functional molecular dynamics simulations, we have demonstrated that ATP binding to the CR3 domain of B-Raf protein induces evolutionarily functionally conserved changes in dynamics that physically connect ATP binding pocket with both the P-loop and activation segment. The region just downstream of the mutation intolerant valine codon 600 is especially affected in its dynamics by ATP binding remotely on the structure of the protein KD. The V600E mutation massively disrupts these evolutionary conserved shifts in dynamics that are obviously a key complex functional feature of BRAF signaling and MAPK regulation.

Hyperactivation inducing drugs, especially the first generation drug sorafenib, appears to mimic functionally conserved dynamics of natural ATP binding function in the CR3 domain, inducing almost the same spatial pattern of functionally conserved protein dynamics in our simulations as ATP itself. Vemurafenib and dabrafenib show less of this tendency, but still induce functional dynamics in the region joining the ATP binding pocket to the P-loop and activation loop segment of B-Raf. In contrast, the hyperactivation breaking drug PLX7904 shows almost no such propensity for mimicry of dynamics of the ATP agonist. Our survey of single amino acid replacements most affecting patient drug sensitivity or resistance to the four inhibitors indicates that mutations impacting the three hyperactivation inducing drugs (i.e. sorefenib, vemurafenib and dabrafenib) appear to target functionally conserved aspects of P-loop and activation loop dynamics. This might be expected under a somatic mutation-selection regime induced by therapeutic treatments of tumors where functional B-Raf dynamics are still potentially in play, albeit likely subverted into a malfunctional state from the standpoint of the patient. However, PLX7904 currently has very few known sensitizing or resistant mutations, an observation that may reflect lack of experimentation with the new compound, but may also prove consistent with our demonstration that it largely avoids inducing the same sort of dynamic changes as ATP upon binding the same location.

Finally, our molecular evolutionary study, indicates that B-Raf sensitivity (i.e. mutational intolerance) to V600E has a likely ancient origin, perhaps as old as the Raf kinase protein family itself. However, the sensitivity of functionally conserved B-Raf dynamics to V600E, appears to have greatly increased sometime during early jawed vertebrate evolution, as represented by extant sequences across all modern ray-finned fishes. As a modulator of multicellular growth, B-Raf likely has origins in early eukaryotes, however poor alignment quality to putative orthologs in *C. elegans* and *S. cerevisiae* prevented us from exploring this possibility further.

## Conclusion

We propose that functional dynamic mimicry (FDM), as observed here in B-Raf inhibitors, is an important risk factor to consider in future rational drug design. This an especially important consideration when targeting sites of natural agonists involved in signal activation in pathways like MAPK which are typically downregulated after early embryonic development, and a subsequently problematic in all cancers. Pharmaceuticals that avoid FDM while effectively targeting and binding active sites of proteins in these pathways, are most likely disrupting the functional dynamics of protein domains to the extent that they are no longer recognized by the pathways in which they normally participate. In the case of pathways downregulated after early development, this affect is ideal, as it destroys signal transduction. Our computational software (Babbitt et al., 2019, 2018b) can allow for inexpensive screening against unintended effects of FDM, potentially producing better patient products while saving pharmaceutical industry wasted time, effort and money in preclinical development and clinical trial. We also conclude that a more evolutionary and biophysical perspective around complex problems like FDM that are potentially linked to naturally evolved tolerances to protein mutation are greatly needed in future biomedical research. This cannot be easily addressed by the combination of DNA sequence-based bioinformatics and cell/animal-based experimental research alone. Proteins are nanometer sized machines operating in the complex context of soft matter dynamics. They are often imperfect in their interactions with other proteins and incompletely optimized in terms of their functional evolution. The development of modern drug therapies could benefit by embracing a more functional evolutionary perspective of putative protein targets in addition the more automated and comprehensive high-throughput exploration of small molecule space currently underway. B-Raf’s apparently ancient intolerance to position 600 mutation along with its clear and singular role in tumor progression and drug resistance might imply that evolution has had ample enough time to explore protein space near this site, with no success in finding a more robust solution to MAPK regulation. But it also implies that perhaps it is time for humanity to consider re-engineering somatic protein targets to better tolerate existing therapeutics and common mutations, rather than to endlessly search through variations of small molecule drugs that are not ever guaranteed to potentiate cures.

## List of abbreviations

ATP: adenosine triphosphate
BRAF: B-Raf gene sequence
B-Raf: protein product of BRAF gene
BRAFi: B-Raf inhibitor
CC: canonical correlation
CR1 CR2 CR3: conserved region 1, 2, or 3
DNA: deoxyribonucleic acid
dFLUX: change in atom fluctuation or rmsf
DROIDS: software acronym for “detecting outlier impacts in dynamic simulation”, our software package for comparative protein dynamics simulation
FDM: functional dynamic mimicry
GPU: graphic processor unit
KD: kinase domain
KL divergence: Kullback-Leibler divergence
MAPK: mitogen-activated protein kinase
maxDemon: software acronym for “Maxwell’s demon”, our machine learning application for DROIDS
MD: molecular dynamic (simulation)
NCBI: National Center for Biotechnology Information
PDB: Protein Data Bank
P-loop: ATP stabilizing loop
rmsf: root mean square fluctuation
V600E: valine to glutamic acid mutation and position 600

## Acknowledgements

We acknowledge Nvidia corporation for a GPU hardware grant, and the College of Science at the Rochester Institute of Technology for internal grant support.

The authors declare no conflicts of interest.

## Appendix A

**Supplementary information on the four inhibitors:** The four B-Raf inhibitors (BRAFi) discussed in this manuscript are all small molecule compounds that are FDA approved for use: 1. in mutated BRAF in metastatic melanoma (Vemurafinib, Dabrafenib) or 2. in other carcinomas (Sorafenib, for advanced renal cell, thyroid, and hepatocellular carcinomas) as a more general kinase inhibitor (active against BRAF, VEGFR, etc). PLX7904 is in preclinical development.

**Sorafenib** (also, Nexavar, BAY 43-9006, MW: 464.8 g/mol): A member of the class of phenylureas that acts as a kinase inhibitor and angiogenesis inhibitor. It is nonspecific, and does not distinguish between WT and mutant B-Raf forms, and has higher specificity for C-Raf compared to B-Raf. It is known to potently induce hyperactivation, and leads to rapid development of drug resistance.

**Vemurafinib** (also, PLX4032; Zelboraf; MW: 489.9 g/mol): An ATP-competitive kinase inhibitor selective for B-Raf V600E. It functions to decrease signaling through the MAPK pathway by inhibiting the kinase domain of mutant BRAF, but is known to also induce hyperactivation in WT BRAF. Shows increased specificity for B-Raf V600E relative to B-Raf WT or C-Raf.

**Dabrafenib** (also, Tafinlar, GSK2118436; 519.6 g/mol): An ATP-competitive kinase inhibitor selective for B-Raf V600 (E/K/R) mutations, which acts to induce inhibition of phosphorylated extracellular signal-regulated kinase (ERK). Often given in combination with MEK inhibitor Trametinib to achieve a more favorable resistance profile.

**PLX7904** (also PB04; MW: 512.5 g/mol): A selective B-Raf kinase inhibitor known to be a potent paradox-breaker. able to efficiently inhibit activation of ERK1/2 in mutant BRAF melanoma cells but does not hyperactivate ERK1/2 in mutant RAS-expressing cells. Was designed to interact with position 505 in B-Raf, shifting the Raf dimer interface, and cause inhibition via long-range structural alterations. Along with analogue PLX8394, is in development as BRAFi capable of uncoupling RAF blocking and MAPK pathway activation.

