## Supplemental Materials for "Function and evolution of B-Raf loop dynamics relevant to cancer recurrence under drug inhibition"

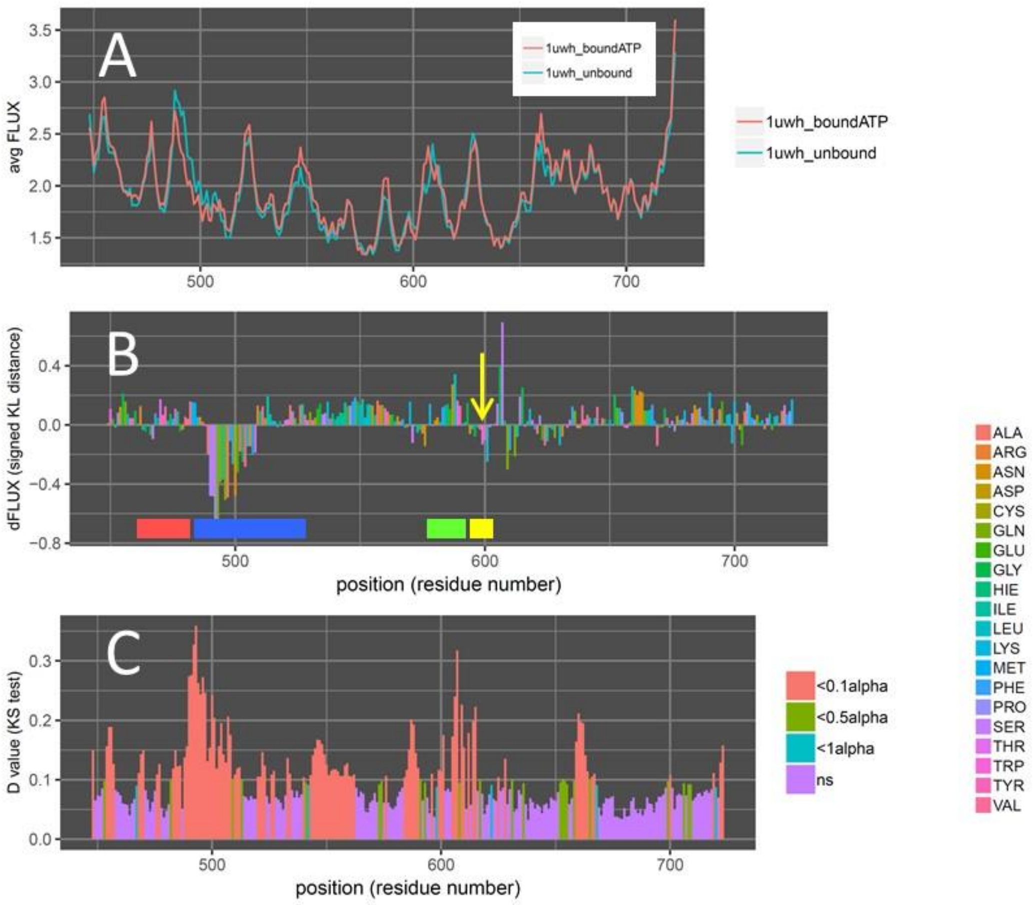


***Supplemental Figure 1. Additional comparative molecular dynamic analyses for ATP interaction in wild type BRAF.*** *(A) Average atom fluctuation over time (i.e. RMSF or FLUX) for two ensembles of 300 x 0.5ns molecular dynamic simulations of inhibitor bound and unbound structures are shown and statistically compared. (B) Signed symmetric Kullback-Leibler (KL) divergences (i.e. relative entropy) in local RMSF distributions (i.e. dFLUX) from each amino acid backbone are plotted with positive values indicating amplified fluctuation and negative values indicating stabilized regions. Colored bars indicate the P-loop (red), ATP binding pocket (blue), catalytic loop (green) and a highly flexible activation loop (yellow). P values from a Benjamini-Hochberg corrected Kolmogorov-Smirnov (KS) test are also shown in (C) indicating significant differences in dynamics between bound and unbound structures.*

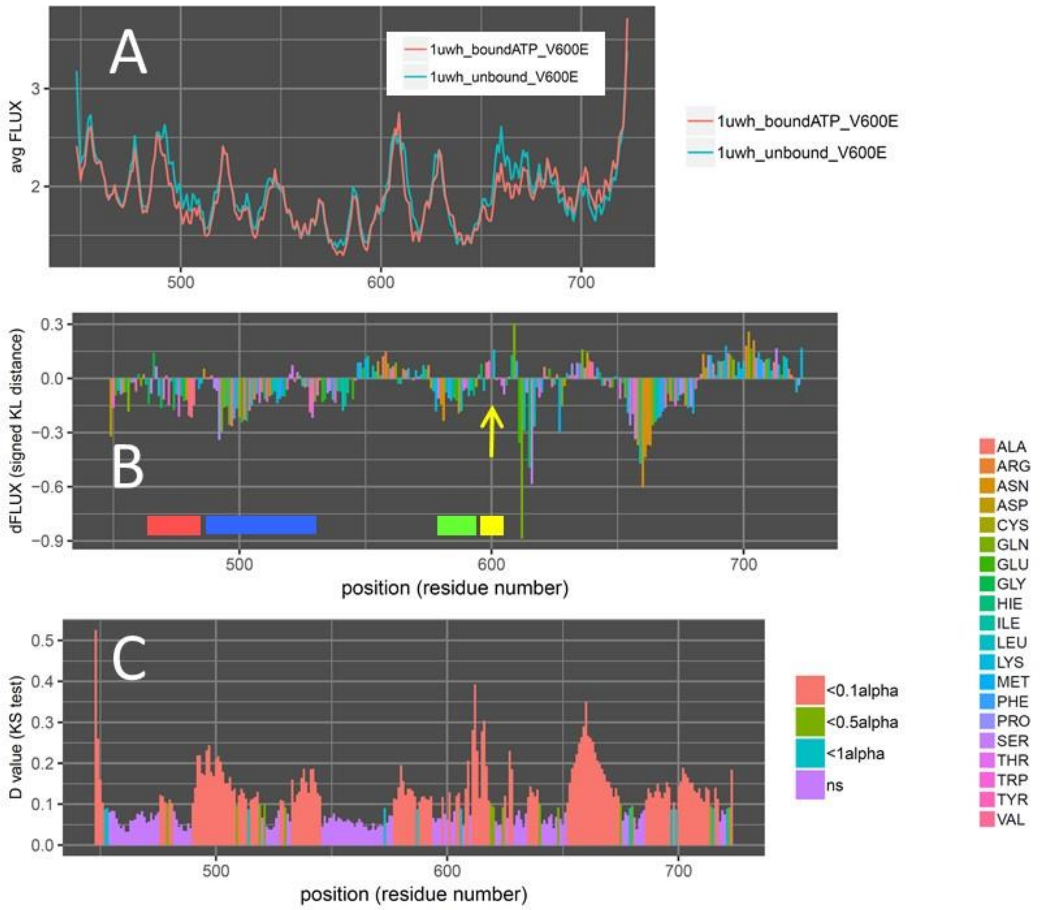


***Supplemental Figure 2. Additional comparative molecular dynamic analyses for ATP interaction in V600E mutant BRAF.*** *(A) Average atom fluctuation over time (i.e. RMSF or FLUX) for two ensembles of 300 x 0.5ns molecular dynamic simulations of inhibitor bound and unbound structures are shown and statistically compared. (B) Signed symmetric Kullback-Leibler (KL) divergences (i.e. relative entropy) in local RMSF distributions (i.e. dFLUX) from each amino acid backbone are plotted with positive values indicating amplified fluctuation and negative values indicating stabilized regions. Colored bars indicate the P-loop (red), ATP binding pocket (blue), catalytic loop (green) and a highly flexible activation loop (yellow). P values from a Benjamini-Hochberg corrected Kolmogorov-Smirnov (KS) test are also shown in (C) indicating significant differences in dynamics between bound and unbound structures.*

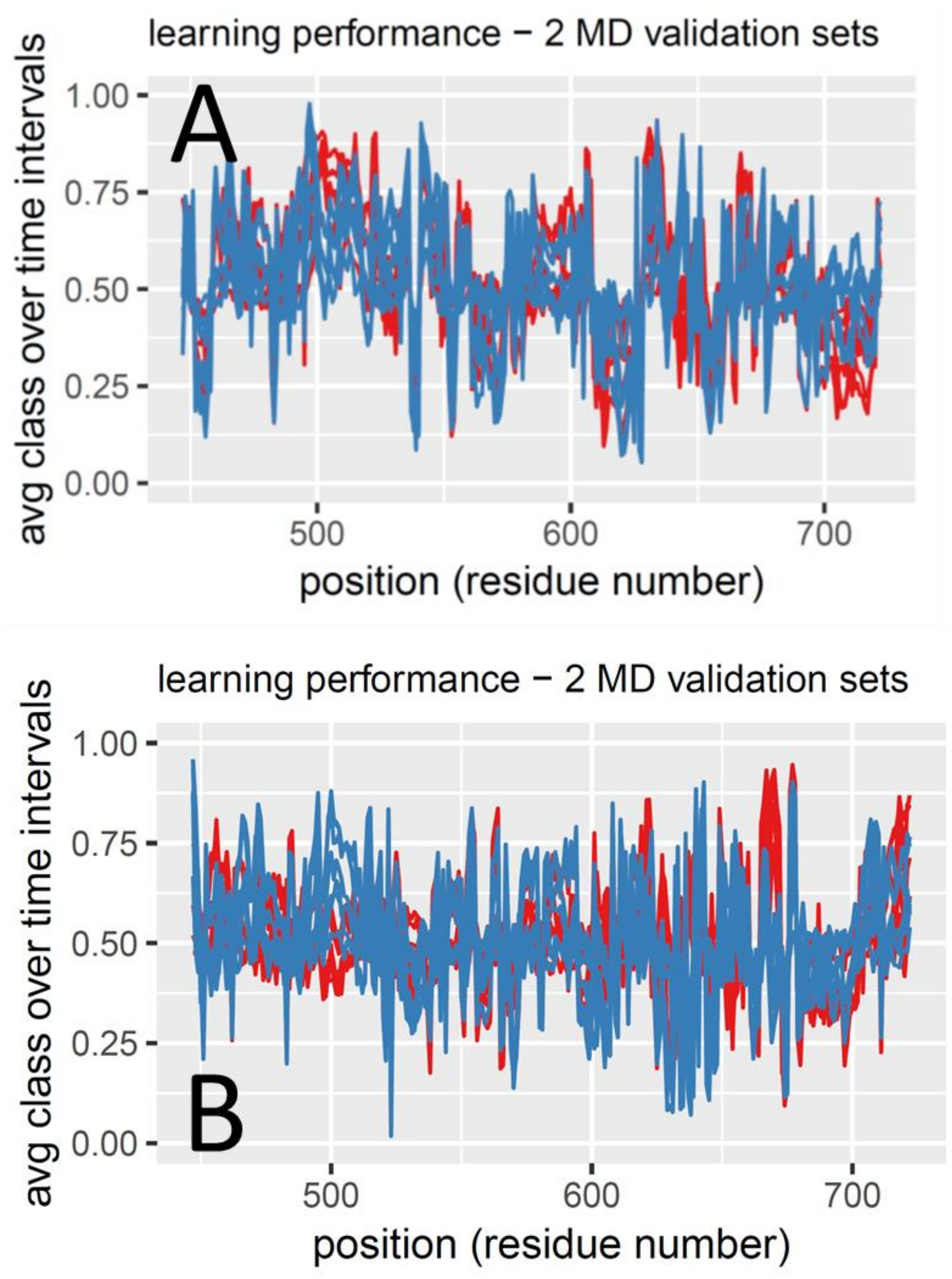


**Supplemental Figure 3. Raw local machine learning performance used to derive canonical correlations for detecting functionally conserved dynamics are shown for comparison in both (A) wildtype and (B) V600E mutant backgrounds. Two validation runs are plotted (red and blue) using all seven machine learners. The greater correlation in A compared to B can be seen, indicating the effect of V600E in destroying the functional dynamics of B-raf that is normally responsive to ATP binding.**

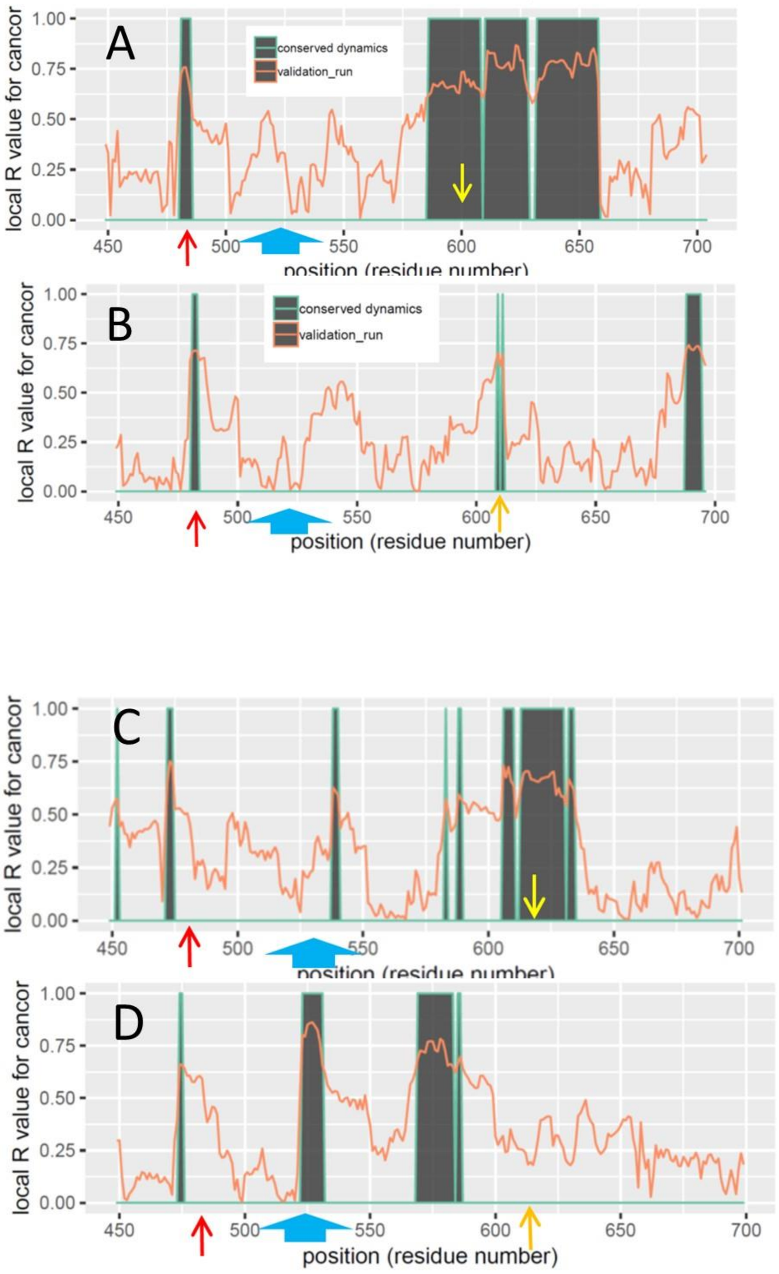


***Supplemental Figure 4. Position plots for the comparisons of the effects of four BRAF inhibitors on functionally conserved dynamics of ATP binding interaction in human BRAF kinase domain.*** *Inhibitors are (A) sorafenib, (B) PLX7904, (C) vemurafenib and (D) dabrafenib.*

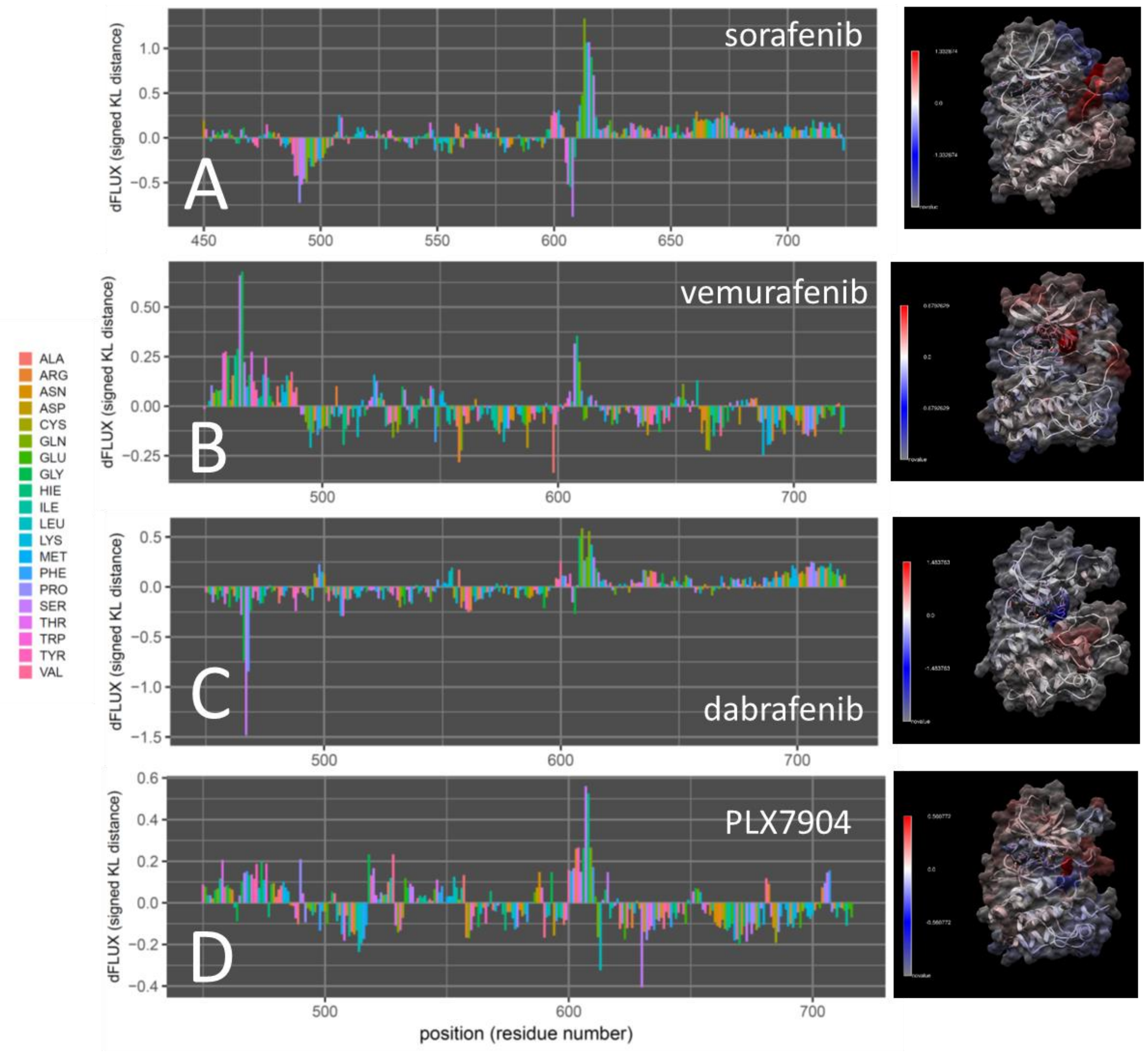


**Supplemental Figure 5. Comparative molecular dynamic analysis of Raf inhibitor binding interactions in normal wild type human B-Raf kinase domain. Inhibitors are (A) sorafenib, (B) vemurafenib, (C) dabrafenib and (D) PLX7904 (PDB ID = 1uwh, 4rzv, 4xv2 and 4xv1). Signed symmetric Kullback-Leibler (KL) divergences in local RMSF distributions (i.e. dFLUX) from each amino acid backbone are plotted (left) shown color mapped on the protein structure (right) with red indicating amplified fluctuation and blue indicating stabilized regions.**

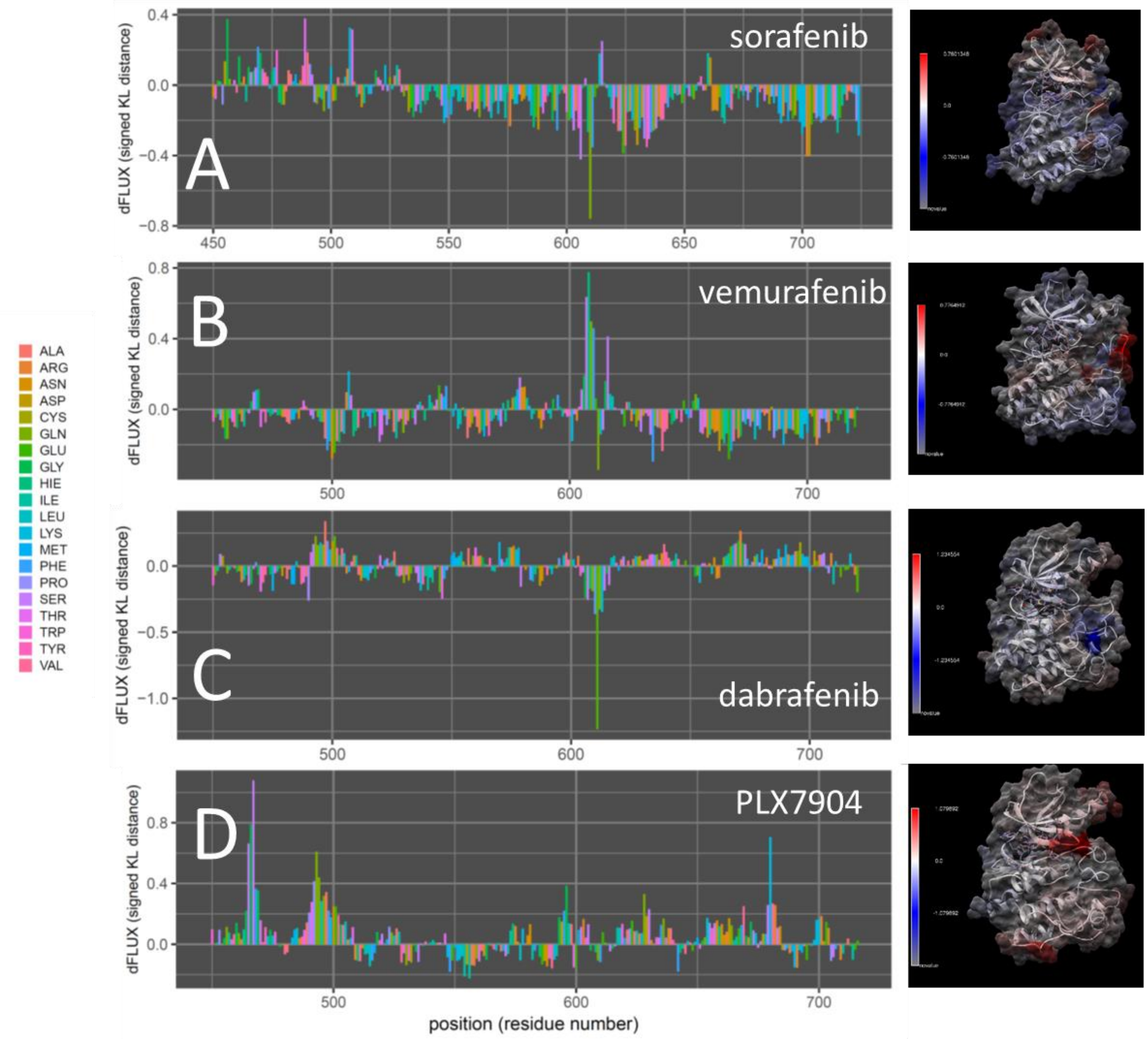


**Supplemental Figure 6. Comparative molecular dynamic analysis of Raf inhibitor binding interactions in mutant V600E human B-Raf kinase domain. Inhibitors are (A) sorafenib, (B) vemurafenib, (C) dabrafenib and (D) PLX7904 (PDB ID = 1uwh, 4rzv, 4xv2 and 4xv1). Signed symmetric Kullback-Leibler (KL) divergences in local RMSF distributions (i.e. dFLUX) from each amino acid backbone are plotted (left) shown color mapped on the protein structure (right) with red indicating amplified fluctuation and blue indicating stabilized regions.**

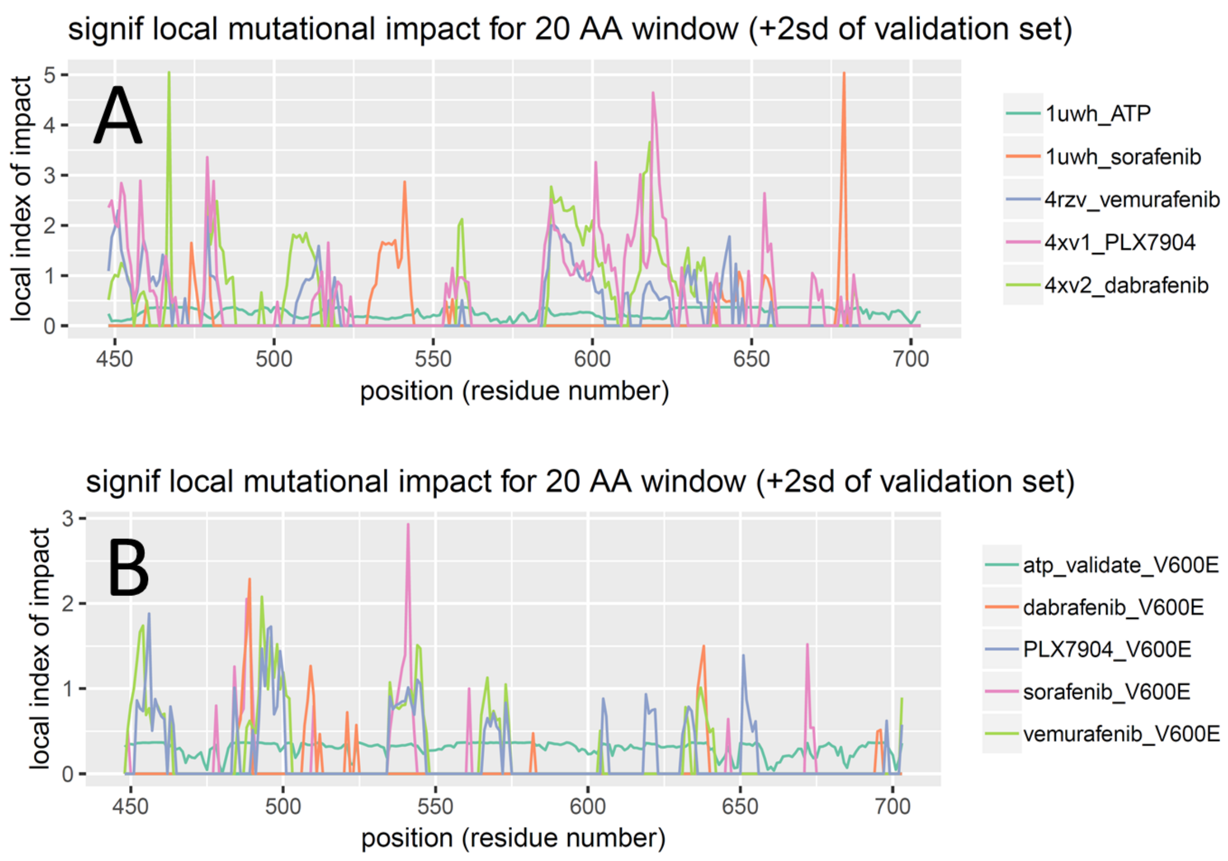


**Supplemental Figure 7. Local impacts of four Raf inhibitor drug class variants on functionally conserved ATP binding dynamics in B-raf protein compared in both (A) wildtype and (B) V600E mutant backgrounds. Additional runs with just ATP binding impact is also shown for validation.**

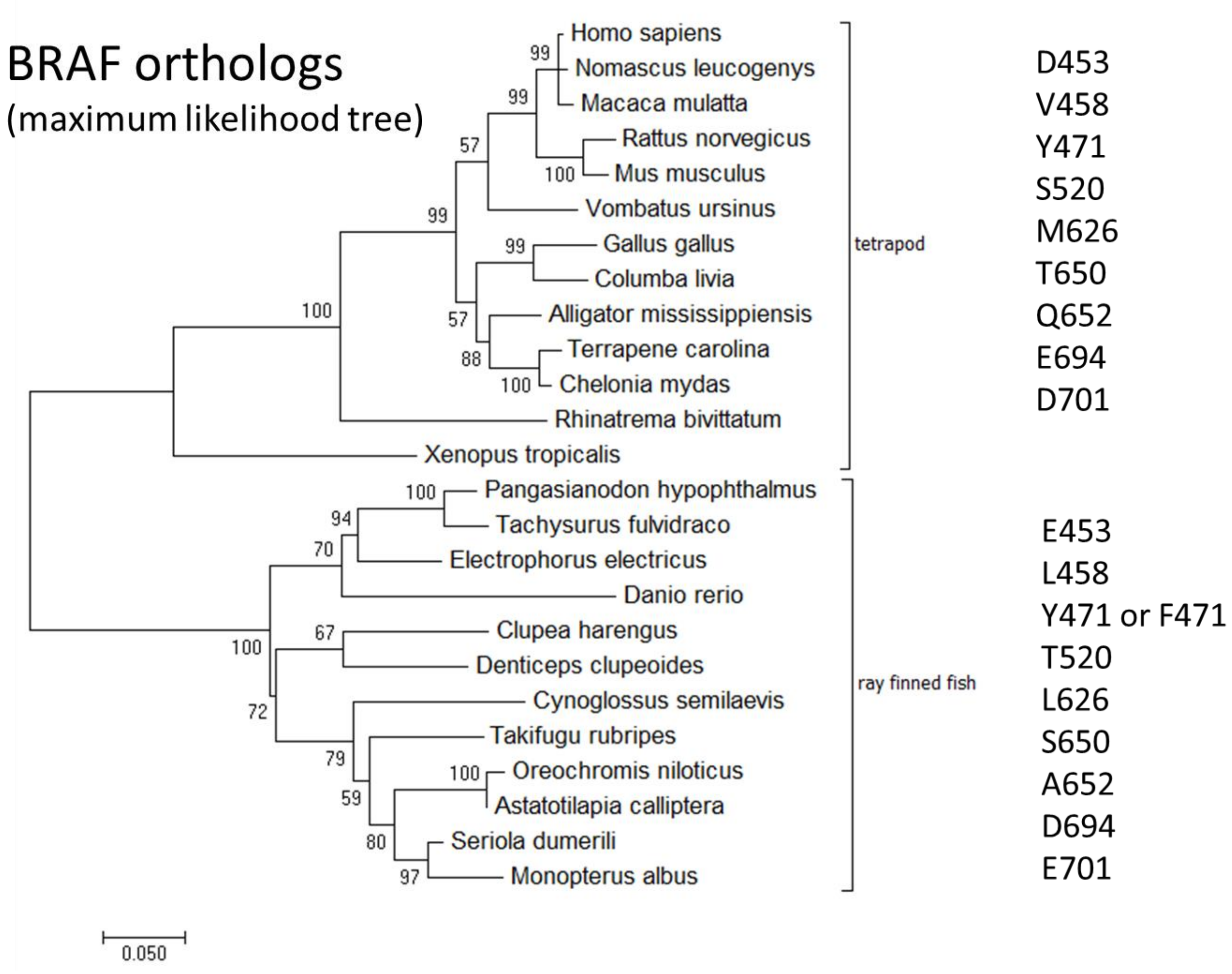


**Supplemental Figure 8. A maximum likelihood tree bootstrapped 500 times using the Kimura 2 parameter model with Gamma correction for multiple hits and invariant sites. This best model was determined via multi-model inference (BIC, AICc). Insect outgroup is not shown to enhance readability of the bootstrap values.**
